# Accurate tiling of spatial single-cell data with Tessera

**DOI:** 10.1101/2025.01.17.633630

**Authors:** Daniel J. Stein, Miles Tran, Ilya Korsunsky

## Abstract

Single-cell spatial transcriptomics reveals how cells organize in healthy and diseased tissues. From these data, tissue segmentation analysis defines discrete compartments that organize cells into functional multicellular units. Existing methods for automated tissue segmentation rely on spatial smoothing to define spatially coherent regions but often blur the boundaries between adjacent tissue compartments. We describe Tessera, an algorithm that approaches tissue segmentation through a novel approach, dividing the tissue into small multicellular tiles whose edges track with natural tissue boundaries. Tessera achieves this by incorporating successful tools from edge-preserving smoothing, topological data analysis, and morphology-aware agglomerative spatial clustering. We show that Tessera identifies a range of known anatomical structures, in healthy mouse brain and human lymph nodes, and novel disease-associated niches, in human brain and in lung cancer. Tessera is a general-purpose tool that returns spatially coherent spatial structures with accurate boundaries across a range of spatial transcriptomics and proteomics technologies.

## Introduction

Tissues are organized into tightly regulated compartments, each with a distinct composition of cells that work together to enact physiological or pathological functions. Our collective knowledge of cellular heterogeneity has grown to define 100s of cell types and 1000s of functional cell states^1,2^. Observing how these cells organize into functional tissue compartments is now feasible, with the advent of commercialized platforms that spatially profile 10s of proteins and 1000s of genes in single cells *in situ*. The major gap to map this organization is analytical. It is challenging to accurately define these tissue compartments, especially in diseased tissues, where boundaries tend to break and shift. Moreover, even when regions are well characterized (e.g. cortical brain layers), it is not trivial to accurately map boundaries with micron-level accuracy. Thus, we need a general-purpose analysis tool to identify distinct tissue regions in an automated, scalable, and accurate way.

Modern algorithms automate tissue segmentation by framing it as spatial clustering, grouping cells by their spatial neighborhood identity rather than by intrinsic cell state. These algorithms maximize coherence (i.e. smoothness) of spatial clusters through spatial smoothing, achieved either by spatial feature aggregation (e.g. UTAG^3^ and CellCharter^4^) or by incorporating spatial location into dimensionality reduction with variants on random Markov fields (e.g. HMRF^5^, STAGATE^6^, and BayesSpace^7^). For both approaches, tuning parameters to achieve the right amount of smoothness is important: too little smoothness diminishes spatial coherence while too much blurs the boundaries between neighboring tissue compartments. Unfortunately, it is impossible to *a priori* set the optimal parameters, especially since the amount of smoothness needed differs from one location to another. This is a well-studied problem in image analysis, with many solutions based on anisotropic (i.e. spatially asymmetric) smoothing^8^ and edge-informed image segmentation^9,10^. The goal of these methods is to extract spatially coherent objects with well-defined edges automatically from potentially noisy 2D images.

In this manuscript, we first describe the high level components of Tessera, our novel algorithm for boundary-sensitive tissue tiling, and then demonstrate these steps on one sample of the mouse brain, with well annotated regions. These steps are wrapped into a convenient function in the Tessera R package. We then demonstrate the accuracy, speed, and utility of Tessera in a set of increasingly complex scenarios. In healthy tissue, Tessera recovers known anatomical structures in a variety of tissues, even when boundaries are thin or noisy. In diseased tissue, Tessera finds pathological structures associated with disease progression and predictive of clinical outcome. Overall, Tessera represents the first general purpose algorithm to successfully extract spatially coherent multicellular regions with accurate boundaries from single-cell spatial genomics data.

## Results

### Tessera groups cells into tiles along natural boundaries

The purpose of Tessera is to group spatially adjacent cells into multicellular tiles, whose boundaries correspond to natural tissue boundaries between anatomical and functional multicellular structures (**Fig 1a**). The Tessera algorithm consists of four steps, described in detail in **Methods**. In the first step (**Fig 1b**), we compress the molecular information (e.g. gene or protein expression) inside cells into a low dimensional representation (by default, with PCA), without using any spatial information. In the second step (**Fig 1c**), we look at each cell’s local spatial neighbors and compute the direction and magnitude of the largest change in PCA embeddings within each neighborhood. The goal here is to distinguish between cells inside a tissue compartment (i.e. low magnitude) versus those at the boundary between compartments (i.e. high magnitude). In practice, these gradients are sensitive to noise. Therefore, we apply a direction-aware smoothing operator—anisotropic bilateral filtering (**Methods**)—to encourage consistency between gradients of physically adjacent cells. In the third step (**Fig 1d**), we trace out boundaries between cells with large spatial gradients using Discrete Morse Theory (**Methods**) and end up with a set of tiles whose edges match these traced boundaries. These tiles are often small, as the traced boundaries are sensitive to small, local changes in gradients. In the fourth step (**Fig 1e**), we aggregate adjacent tiles that have similar molecular profiles (i.e. PCA embeddings) and form reasonable shapes, as quantified with multiple morphological measures, such as area and convexity. This results in the final output of Tessera: larger tiles that represent whole or pieces of real multicellular spatial structures in tissue. These tiles can then be used in downstream analysis to label and define tissue regions.

**Figure 1.**
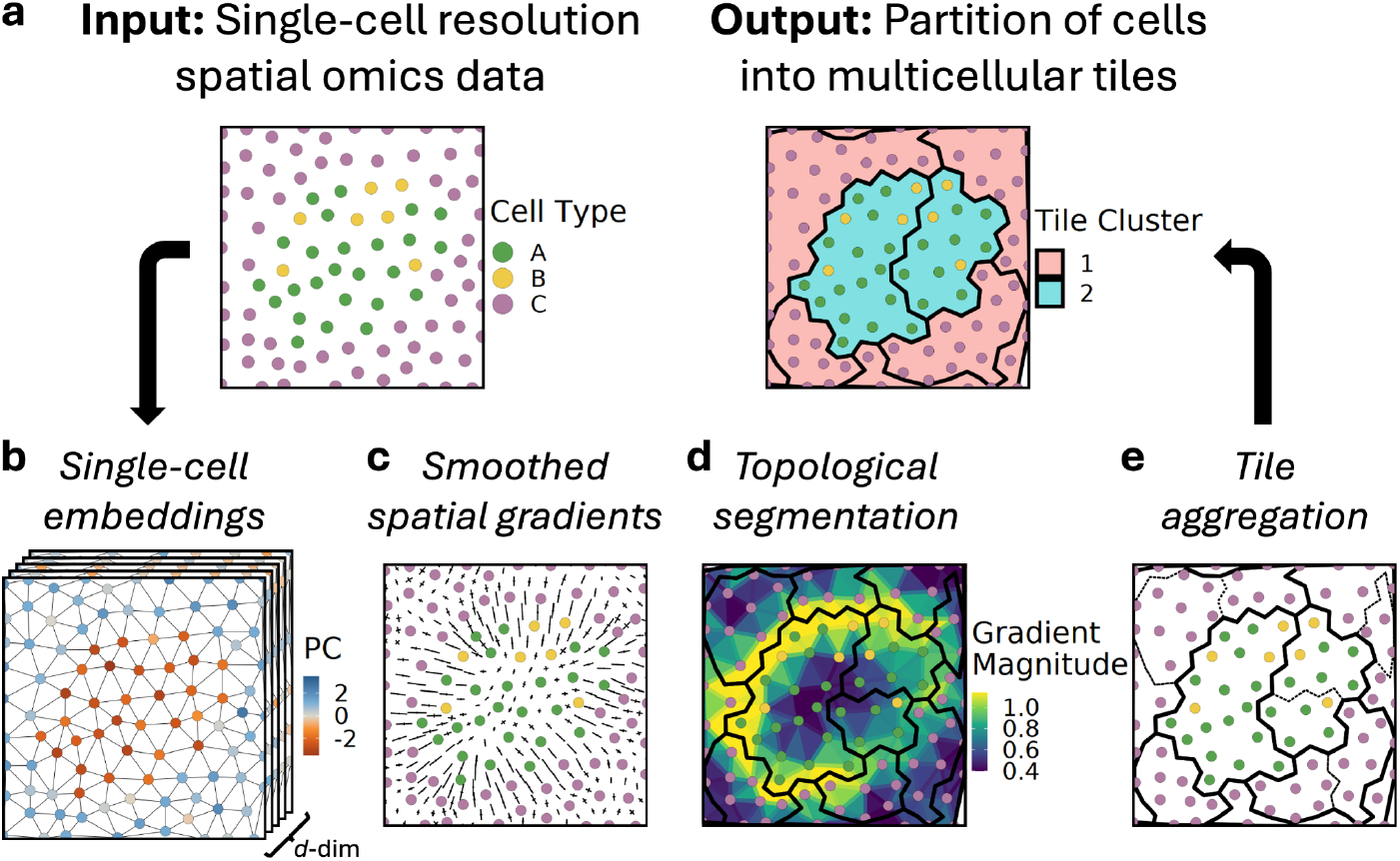
Tiling of spatial single-cell data with Tessera. Overview of the Tessera method. **a**, Each point represents a single cell profiled using spatial transcriptomics or spatial proteomics. Only expression measurements and spatial coordinates are used by Tessera to create and cluster tiles—cell annotations are not used as input. **b**, Single-cell PCA embeddings are computed without using spatial information. Each cell is represented by a *d*-dimensional embedding. **c**, Spatial gradients are computed across all embedding directions and smoothed using anisotropic bilateral filtering. **d**, The gradient magnitude field is segmented using discrete Morse theory. **e**, Initial tiles are aggregated hierarchically using similarity in cell embeddings, number of cells per tile, and compactness of tile shape.

### Accurate domain detection in mouse brain

To demonstrate Tessera in a real dataset, we use Tessera tiles to identify well-characterized anatomical regions in the mouse brain. For exposition, we analyzed one coronal section, covering the cerebral cortex, hippocampus, thalamus, and hypothalamus of one mouse brain, profiled using a 1,122-gene MERFISH panel^11^. In this section, we first walk through the four steps of Tessera and visualize intermediate results on a small 6.25mm^2^ inset of the full sample. Then, we show the final Tessera output, with downstream region annotations, on the full sample.

In step one, we projected the expression of 1,122 genes in 46,339 cells into 50 PCs. While these PCs are defined with only cell-intrinsic information, they also implicitly reflect the spatial distribution of cells in the brain (**Fig 2a**). Here, PC1 separates neuronal vs non-neuronal cells, while PC40 identifies glutamatergic neurons in the CA3 field of the hippocampus (**Fig S1a,b)**. In step two, we define spatial-transcriptional gradients for each cell using the 50 PCs computed in step one. As expected, these gradients are initially noisy (**Fig 2b**). Our anisotropic filter (**Methods**) smooths out these gradients, whose orthogonal components clearly show edges between tissue regions (**Fig 2c**). In step three, we use the vector field to estimate something akin to a geological topographic map (**Fig 2d**), with yellow “hills” along areas of large gradients and purple “valleys” in homogeneous regions. We then use Discrete Morse Theory (**Methods, Fig S2**) to trace out tiles along this map (**Fig 2e**). In the full sample, we get 5,573 small tiles, with an average of 8.3 cells per tile (**Fig S1c)**. In step four, we aggregate adjacent tiles with similar cell type compositions to form more spatially coherent, larger tiles (**Fig 2f)**. In total, we recover 1,384 tiles, with an average of 33 cells per tile (**Fig S1d**).

**Figure 2.**
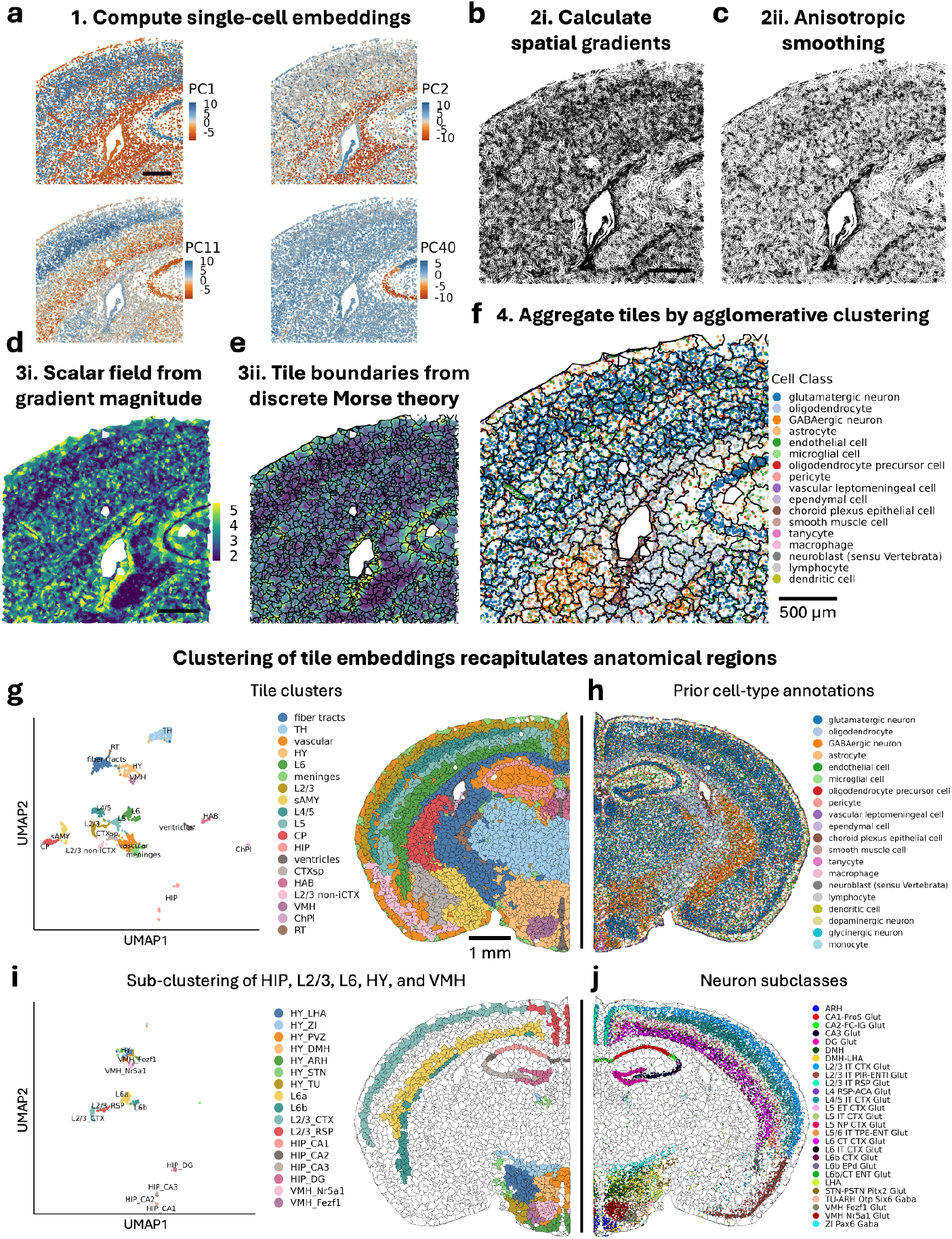
Accurate domain detection in a single section of the mouse brain. Tessera was applied to one coronal section of the mouse brain, profiled with a 1,122-gene MERFISH panel^16^. **a-f**, Step-by-step Tessera results are shown for a 2.5×2.5mm subsection of the sample (scale bar = 500µm). **a**, Single-cell embeddings computed by PCA, with four of 50 total dimensions shown. Each point is an individual cell. **b**, Spatial gradients are computed across all embedding directions. The displayed lines are orthogonal to the principal gradient direction, with length proportional to the gradient magnitude. **c**, Orthogonals to the spatial gradient after anisotropic bilateral filtering. **d**, Magnitude of the total gradient. **e**, Tile boundaries from topological segmentation, along with the underlying gradient magnitude (see **Fig S2** for details). **f**, Tile boundaries after agglomerative clustering of initial tiles from (**e**), where merging is prioritized by similarity in cell embeddings, number of cells per tile, and compactness of tile shape. Each point is an individual cell, colored by prior cell type annotations, which are not used by Tessera. **g-j**, Results from clustering Tessera tiles by expression using modularity-based Leiden clustering (scale bar = 1mm). **g**, UMAP (each point represents a tile) and spatial localization of tile clusters, produced by clustering all 1,384 tiles by their average cell embeddings. **h**, Cells from the same section colored by previous cell type annotations. Coordinates are reflected horizontally for visualization purposes. **i**, UMAP and spatial localization of tile sub-clusters, produced by separately clustering tiles from each of the HIP, L2/3, L6, HY, and VMH regions in (**g**). **j**, Spatial localization of selected neuron subclasses. Abbreviations used in (**g**,**i**): TH, thalamus; HY, hypothalamus; sAMY, striatum-like amygdalar nuclei; CP, caudoputamen; HIP, hippocampus; CTXsp, cortical subplate; HAB, habenula; non-iCTX, non-isocortex; VMH, ventromedial hypothalamic nucleus; ChPl, choroid plexus; RT, reticular nucleus of the thalamus; LHA, lateral hypothalamic area; ZI, zona incerta; PVZ, periventricular zone; ARH, arcuate hypothalamic nucleus; STN, subthalamic nucleus; TU, tuberal nucleus; DG, dentate gyrus.

Next, we used the Tessera tiles to define distinct regions in the mouse brain. Our approach is similar to analysis of single-cell genomics data (**Methods**). We grouped tiles into 19 distinct clusters of mouse brain regions and visualized the tile clusters in UMAP space (**Fig 2g**). These tile clusters are also spatially coherent and correspond to meaningful anatomical regions (**Fig 2g**). We found regions with a variety of morphologies, including cortical layers (L2-6) and white matter fiber tracts; finer structures such as vasculature, ventricular ependyma, and the choroid plexus (ChPl); broad regions such as the thalamus (TH) and hypothalamus (HY); and neuroanatomical domains such as the ventromedial hypothalamic nucleus (VMH) (**Fig 2g**). For each anatomical region, we verified the accuracy of our labels by quantifying representative cell type labels enriched in each region (**Fig 2h, S1e,f**). Because we did not use these cell type labels in learning tiles or tile clusters, they serve as a validation of our regions.

We next asked whether further division of tile clusters would reveal more detailed anatomical structures. We subclustered tiles within the L2/3 and L6 layers, the hippocampus, and the hypothalamus (**Fig 2i**). Tiles within each of the subclusters matched the localization of neuron subclasses specific to particular cortical layers, areas of the hippocampus, distinct nuclei within the hypothalamus, and zones within the VMH with different glutamatergic neurons (Nr5a1 and Fezf1) (**Fig 2j, S1e,f**). Overall, Tessera readily identifies previously defined anatomical regions within the mouse brain, across a variety of shapes and scales.

### Comparison to other methods

Next, we compared the performance of other published methods for domain detection on the mouse brain data from **Fig 2**. We chose two representative, well-reviewed methods^12–14^: STAGATE^6^, an auto-encoder framework that uses a graph attention architecture to learn spatially-informed cell embeddings, and CellCharter^4^, which clusters cells based on their neighborhood gene compositions. Using the regions identified in **Fig 2**, we created two benchmarking tasks: 1) cortical layer segmentation to measure performance on large, regular shapes (**Fig 3a-d**); and 2) ventricle detection to identify smaller regions with thin, precise boundaries (**Fig 3e-g**). For each task, we manually labeled regions based on independent cell annotations (**Methods, Fig S3a**). In both tasks, accurate domain detection will achieve two outcomes: accurate cell-to-domain assignment and spatial coherence. For assignment accuracy, we used the F1 statistic, ranging from 0 (bad) to 1 (good) (**Methods**). For the latter, we quantified a spatial incoherence score as the average fraction of each cell’s neighbors not assigned to the same spatial domain (**Methods**). Here, 0 is perfect coherence, while 1 implies that every pair of neighbors are assigned to different domains. In each task, we ran CellCharter and STAGATE with multiple parameter choices and only reported the most accurate result (**Methods, Fig S3b, S4a**).

**Figure 3.**
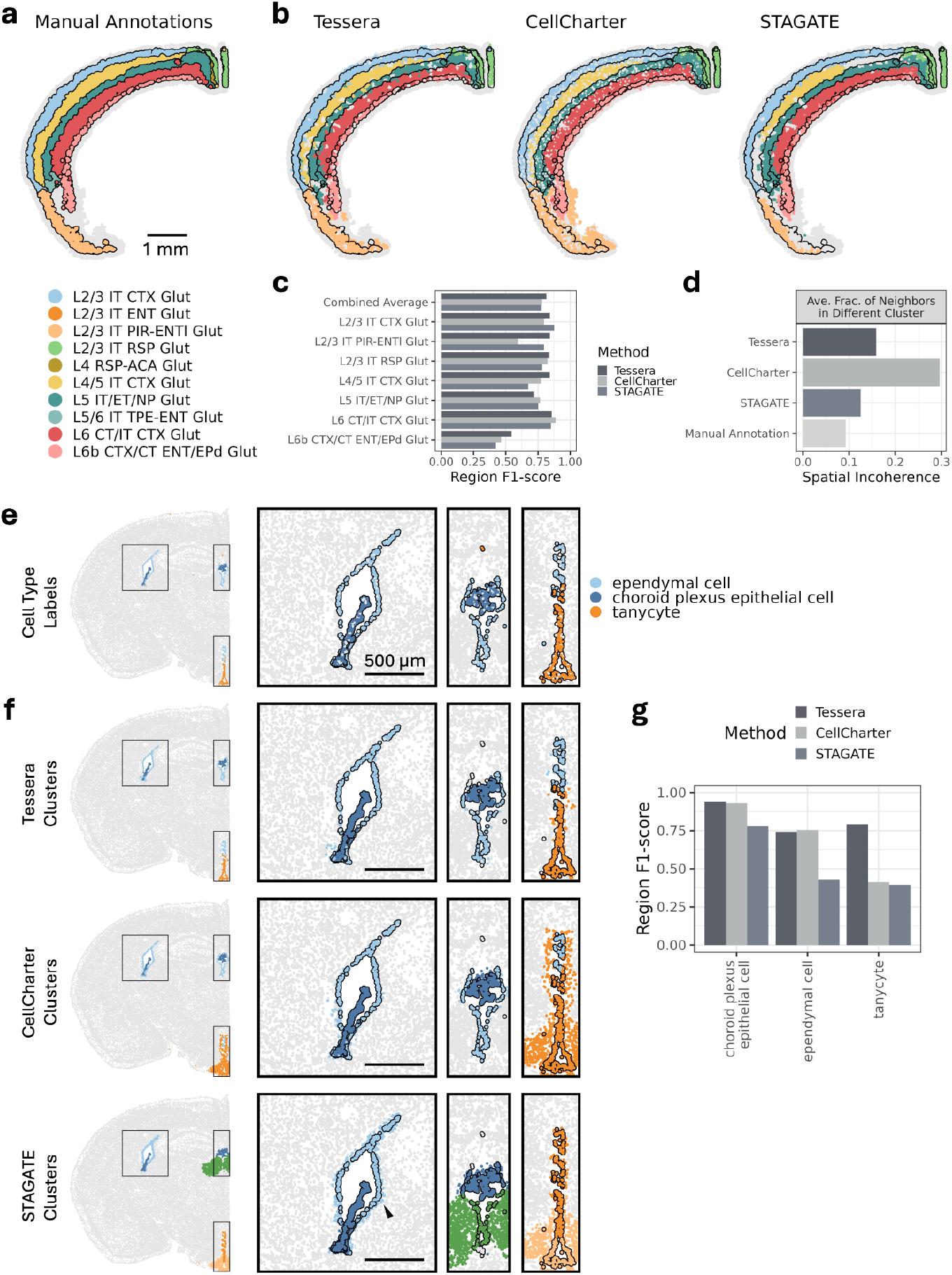
Tessera improves delineation of fine anatomical structures within the mouse brain while maintaining spatially coherent domains. **a-d**, Benchmarking on cortical regions. **a**, Manual annotation of cortical regions based on the distribution of layer-specific glutamatergic neurons. **b**, Spatial domains identified using the three methods Tessera, CellCharter, and STAGATE, where the best clustering resolution across a range of values is shown for each (Leiden resolution = 2, GMM k = 28, and Leiden resolution = 1, respectively). **c**, F1-score (between 0 and 1) for identifying each layer using the best matching domain from each method. The mean F1 across all regions is also shown (combined average, weighted by the size of each region). **d**, Spatial incoherence (higher scores correspond to greater mixing of spatial domains) is computed within the cortical regions in (**a**) and (**b**) as the fraction of each cell’s neighbors that are assigned to a differing spatial domain. **e-f**, Benchmarking on ventricular structures. **e**, Spatial localization of three ventricular cell types, previously annotated, which define different ventricular regions—choroid plexus, ependyma, and tanycyte (a specialized ependymal cell). Insets are shown for three separate ventricular structures. **f**, Spatial domains identified using Tessera, CellCharter, and STAGATE, where only the clusters that best match the ventricular regions defined in (**e**) are shown. The best clustering resolution across a range of values is shown for each (Tessera, Leiden resolution = 7.5; CellCharter, GMM k = 42; and STAGATE, Leiden resolution = 0.3). Arrowhead indicates an ependymal layer that STAGATE inflates beyond the true boundary. **g**, F1-score (between 0 and 1) for identifying each ventricular region in (**e**) using the best matching domain from each method in (**f**).

In task 1 (**Fig 3a**), all methods achieved comparable accuracy, assigning cells to their correct cortical layers (**Fig 3b**) with F1 scores above 0.75 for each method in most layers, with a few exceptions: L2/3 IT PIR-ENTl for CellCharter (F1=0.59), L4/5 IT CTX for STAGATE (F1=0.67), and L5 IT/ET/NP for Tessera (F1=0.72) (**Fig 3c**). The thinnest layer, L6b, was the most difficult, and Tessera slightly outperformed the others, with an F1 score of 0.55 vs 0.47 (CellCharter) and 0.42 (STAGATE). Both CellCharter and STAGATE blurred the edges between L6b cortical cells and their neighboring subcortical cells (**Fig S3c**). For spatial incoherence, the differences were larger (**Fig 3d**). Compared to the best possible result (0.09 for ground truth), Tessera (incoherence=0.16) and STAGATE (incoherence=0.13) achieved similar spatial coherence. Additionally, holes within the Tessera and STAGATE domains (**Fig 3b**) were often meaningful multicellular units, such as larger blood vessels with multiple endothelial cells (**Fig S3d**). In contrast, CellCharter assigned an average of 30% of neighbors to discordant clusters (**Fig 3d**). Further analysis showed that CellCharter assigned many cells by cell type rather than by spatial neighborhood, resulting in large numbers of singletons (8% for CellCharter vs <1% for others, **Fig S3e**) and “holey” domains (**Fig 3b**).

In task 2, the ventricle detection analysis (**Fig 3e**), the differences among methods were more pronounced (**Fig 3f**). Tessera achieved high accuracy (F1>0.74) and low spatial incoherence (0.083) for all three ventricular regions (**Fig 3g**). In contrast, STAGATE achieved F1<0.43 on two regions and CellCharter achieved F1<0.42 on one region (**Fig 3g**). As a result, CellCharter assigned 727 cells incorrectly to the 263-cell tanycyte compartment (precision=0.26, recall=0.98) and STAGATE assigned 212 cells incorrectly to the 263-cell tanycyte compartment (precision=0.36, recall=0.44) (**Fig S4c-e**). Even for the ependymal layer that STAGATE identifies correctly (**Fig 3f, arrowhead**), boundaries are inflated, containing cells beyond the true ependymal boundary and producing low precision (0.44, **Fig S4d**). Together, the results of both benchmarks are consistent with the expectation that CellCharter and STAGATE, which—unlike Tessera—both perform spatial smoothing without boundary detection, blur boundaries between regions and cause merging or overestimation of the size of thin structures, such as the L6b cortical layer and brain ventricle structures. Moreover, CellCharter often underperforms in spatial coherence, clustering cells by cell type rather than spatial neighborhood. Tessera, which clearly delineates tissue boundaries when pooling neighboring cells, avoids both of these issues.

### Reproducibility and scale across full mouse brain

We next asked whether Tessera could scale to a multi-sample cohort, robustly identifying structures that recur across multiple samples. We chose to analyze a dataset with consecutive, serially sectioned coronal brain sections from a single mouse. For successfully annotated regions, we expected the composition of labeled regions in adjacent slices to look similar and change smoothly from slice to slice. We ran Tessera on 4.2 million cells from 147 sections, profiled with a 1,122-gene MERFISH panel^11^, and recovered 126,253 tiles, with 33±14 (mean±s.d.) cells per tile (**Fig S5a**). After clustering the tiles from all sections together, we identified 36 major regions, whose locations and cell type compositions correspond well with known anatomical domains annotated in the Allen Brain Atlas^15^ (**Fig 4a,b, S5b-d**). Moreover, the relative composition of each slice was similar to that of its neighbors along the anterior-posterior axis, resulting in a smoothly changing map of compositional changes from the front of the brain to the back (**Fig 4b**). In addition to composition, the reproducible Tessera regions let us trace changes in the shape of important structures, such as the dense pyramidal cell layers and dentate gyrus region within the hippocampus (**Fig 4c**), multiple layers within the olfactory bulb, cerebrum, and cerebellum (**Fig S6a**), white matter tracts spanning the brain (**Fig S6b**), nuclei with specific subclasses of neurons (**Fig S6c**), and the ventricular system (**Fig S6d**).

**Figure 4.**
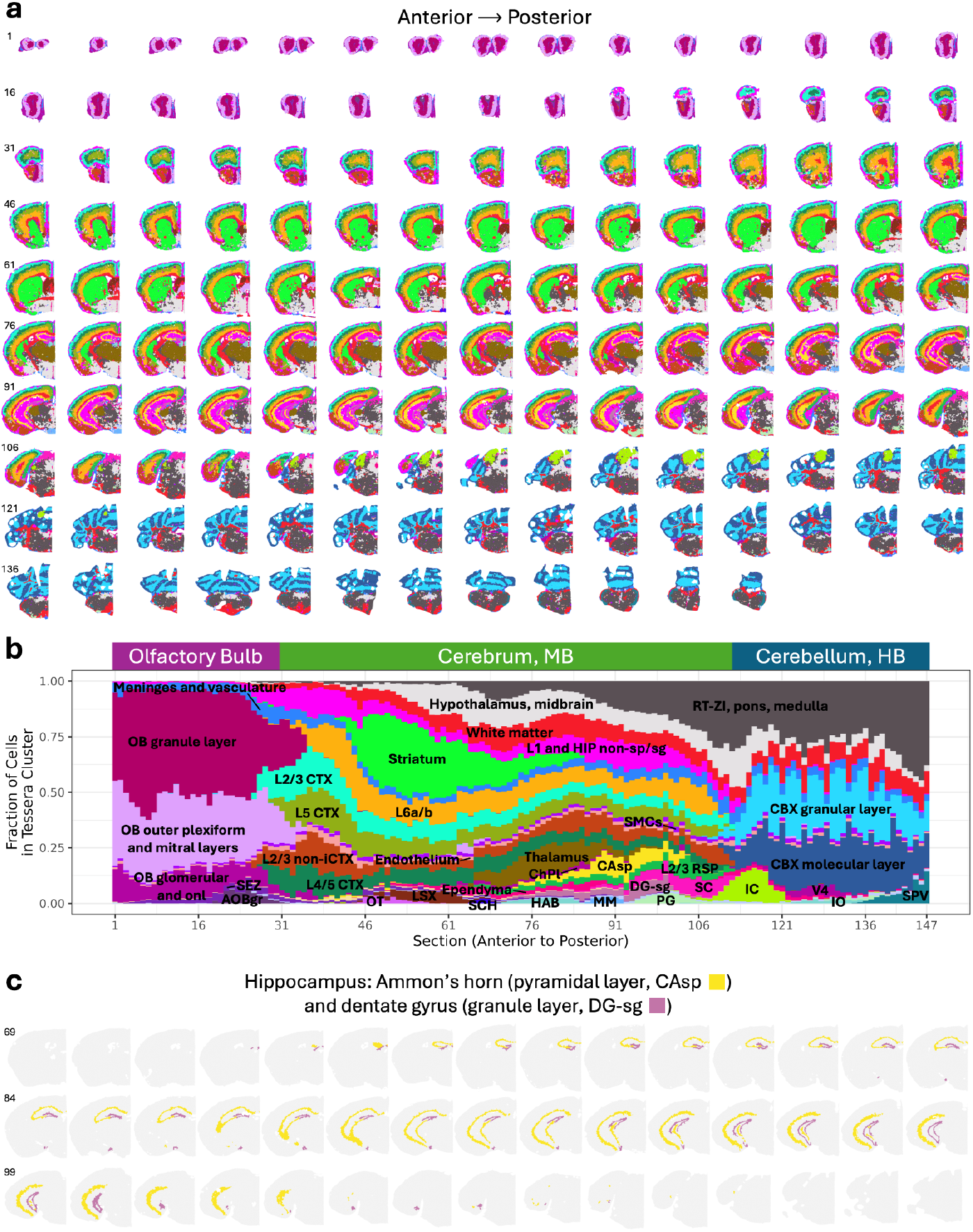
Tessera identifies reproducible anatomical regions across a whole mouse brain. Tessera was applied to 147 consecutive coronal sections of a mouse brain, profiled using a 1,122-gene MERFISH panel, yielding 4.2 million cells^16^. Tessera produced 126,253 tiles, whose embeddings were clustered using modularity-based Leiden clustering. **a**, Spatial localization of tile clusters across all 147 coronal sections of the mouse brain. Sections are ordered from anterior to posterior. Clusters are annotated in (**b**). **b**, Tile cluster composition for each section, ordered from anterior to posterior, and labeled using anatomical regions in the Allen Brain Atlas^15^ (https://atlas.brain-map.org/atlas?atlas=1). The colors of tile clusters match (**a**). **c**, Spatial localization of two hippocampus-associated tile clusters (CAsp and DG-sg) across 45 coronal sections that span the hippocampus. All other tile clusters are shown in grey. Abbreviations used in (**b**): OB, olfactory bulb; MB, midbrain; HB, hindbrain; onl, olfactory nerve layer; AOBgr, accessory olfactory bulb, granular layer; SEZ, subependymal zone; CTX, isocortex; L2/3 non-iCTX, PIR-ENTl-COA-NLOT non-isocortex; OT, olfactory tubercle; LSX, lateral septal complex; SCH, suprachiasmatic nucleus; HIP non-sp/sg, hippocampus non-pyramidal and granule cell layers; ChPl, choroid plexus; HAB, habenula; MM, medial mammillary nucleus; CAsp, hippocampus field CA1/2/3 pyramidal layer; DG-sg, hippocampus dentate gyrus granule cell layer; SMCs, smooth muscle cells (vasculature); PG, pontine gray; RSP, retrosplenial area; RT-ZI, reticular nucleus of the thalamus and zona incerta; SC, superior colliculus; IC, inferior colliculus; CBX, cerebellar cortex; V4, fourth ventricle; IO, inferior olivary complex; SPV, Spinal nucleus of the trigeminal.

Given the large size of this dataset, we took the opportunity to assess the computational resources needed to run Tessera, CellCharter, and STAGATE. We ran each algorithm on 2, 5, 10, 20, 50, 100, and 147 brain slices, corresponding to between 72,000 and 4.2 million cells, to investigate how the algorithms scale with increasing numbers of cells. A fair comparison among these methods is difficult, as each algorithm performs different steps to turn cells into annotated tissue regions. STAGATE performs all steps in one, and CellCharter takes input from the stand-alone scVI software, while Tessera tiling follows PCA (and Harmony) and precedes clustering. We therefore recorded runtimes for CellCharter and Tessera with and without the “additional” steps. Finally, we used the optimal hardware for each algorithm: multi-core CPU for Tessera and GPU for CellCharter and STAGATE (**Methods**). Without scVI or PCA, Tessera (on CPU) processed the full 4.2 million cells faster than did CellCharter (on GPU), requiring only 7.5 minutes (2.5 min to construct tiles + 5 min for Leiden clustering) vs 23 minutes (CellCharter feature aggregation + GMM clustering) (**Fig S5e**). In contrast, STAGATE failed to run on more than one slice, running out of memory before completion. When we include dimensionality reduction and batch correction, CellCharter+scVI processed the full dataset faster, in 34 minutes, than Tessera+PCA+Harmony, which finished in 84 minutes. This difference was driven primarily by scVI, which is slower than PCA+Harmony on smaller datasets (<1.5M cells) but compensates for large data (>1.5M cells) by executing fewer rounds of model optimization. We next broke down the run time for Tessera and its accompanying steps (**Fig S5f**) and found that learning Tessera tiles is faster than Leiden clustering, PCA, or Harmony. Furthermore, Tessera required only 23GB of RAM to process the full 4.2 million cells. Together, these results show that Tessera scales computationally to large multi-sample spatial transcriptomics cohorts without the need for specialized hardware.

### Disease-associated niches in early Alzheimer’s disease progression

We next hypothesized that by finding shared structures across tissue sections, Tessera could enable case-control comparisons to identify anatomical regions associated with treatment or disease severity. We chose the SEA-AD MERFISH dataset, a 140-gene brain atlas of 75 middle temporal gyrus (MTG) samples from 27 donors across a spectrum of Alzheimer’s disease (AD) progression^16^. Each donor had previously been assigned a quantitative disease severity score, called the Continuous Pseudo-progression Score (CPS) between 0 and 1, ranking donors from early disease with early amyloid pathology and some neuronal loss (0-0.4 CPS) to more advanced disease with significant cognitive decline (0.4-1 CPS). Importantly, CPS quantitatively ranks patients within each category^16^. We sought to use Tessera tiles to find brain regions that expanded or contracted in patients with increasing disease pathology, as quantified by CPS.

We applied our standard multi-sample Tessera workflow (**Methods**) to group 1.7M high-quality cells (≥30 transcripts and ≥15 unique genes) into 149K high quality tiles (mean 12±6SD cells per tile, **Fig S7a**), clustered into 17 shared brain regions (**Fig 5a,b, S7b**): seven cortical layers enriched in L2-L6 neurons (**Fig 5b,c, S7c**), seven subcortical regions (i.e. white matter) enriched in oligodendrocytes (**Fig 5b,c**), and four regions marked by vasculature, astrocytes, OPCs, and GABAergic neurons spread throughout the cortex and white matter (**Fig 5b,c, S7b**). In contrast to the regular spatial patterns of the cortical layers, the seven white matter regions were spatially mixed and reflected small pockets of non-neuronal cell types, including microglia, OPCs, and vasculature (**Fig 5c, S7b**). We next asked whether any of these regions expands or diminishes with disease progression. Our approach was to correlate the abundance of each region with donors’ CPS (**Methods**), first across the whole cohort and then separately within early-stage (CPS≤0.4) and late-stage (CPS>0.4) disease, which may have distinct cellular etiologies. We were careful to perform this association separately within the cortex and white matter (**Fig 5d**), as the proportion of cortex to subcortex in each sample varies randomly (**Fig S7d-f**) and can lead to spurious associations. After correcting for multiple hypothesis testing, we found two significant associations (**Fig 5d**), both with early-stage disease progression: decreasing abundance of white matter with microglia-PVM (FDR=0.029, **Fig 5e-g**) and increasing abundance of white matter with GABAergic neurons (FDR=0.029, **Fig S7g**). These results suggest that the decrease of some microglia or immune subtype in the white matter may be important to early disease progression. To resolve the exact subtype, we performed deep annotation of the microglia-PVM cells into 7 classes, previously defined by Gabitto et al^16^. Within the microglia-PVM-enriched white matter tiles, we found Micro-PVM_3-SEAAD microglia (*ZNF804A*^+^) to be the most significantly enriched state (**Fig 5h**). Gabitto et al^16^ found these same Micro-PVM_3-SEAAD microglia expanded in late-stage (CPS>0.6) patients in an analysis of scRNA-seq data of the SEA-AD cohort. However, because Micro-PVM_3-SEAAD microglia are present in both cortex and white matter (**Fig S7h**) and scRNA-seq captured cells from both compartments, it is impossible to know whether the scRNA-seq association is driven by immune cells in the cortex or white matter. By identifying shared tissue regions across samples, Tessera enables direct comparisons of niches within these distinct compartments.

**Figure 5.**
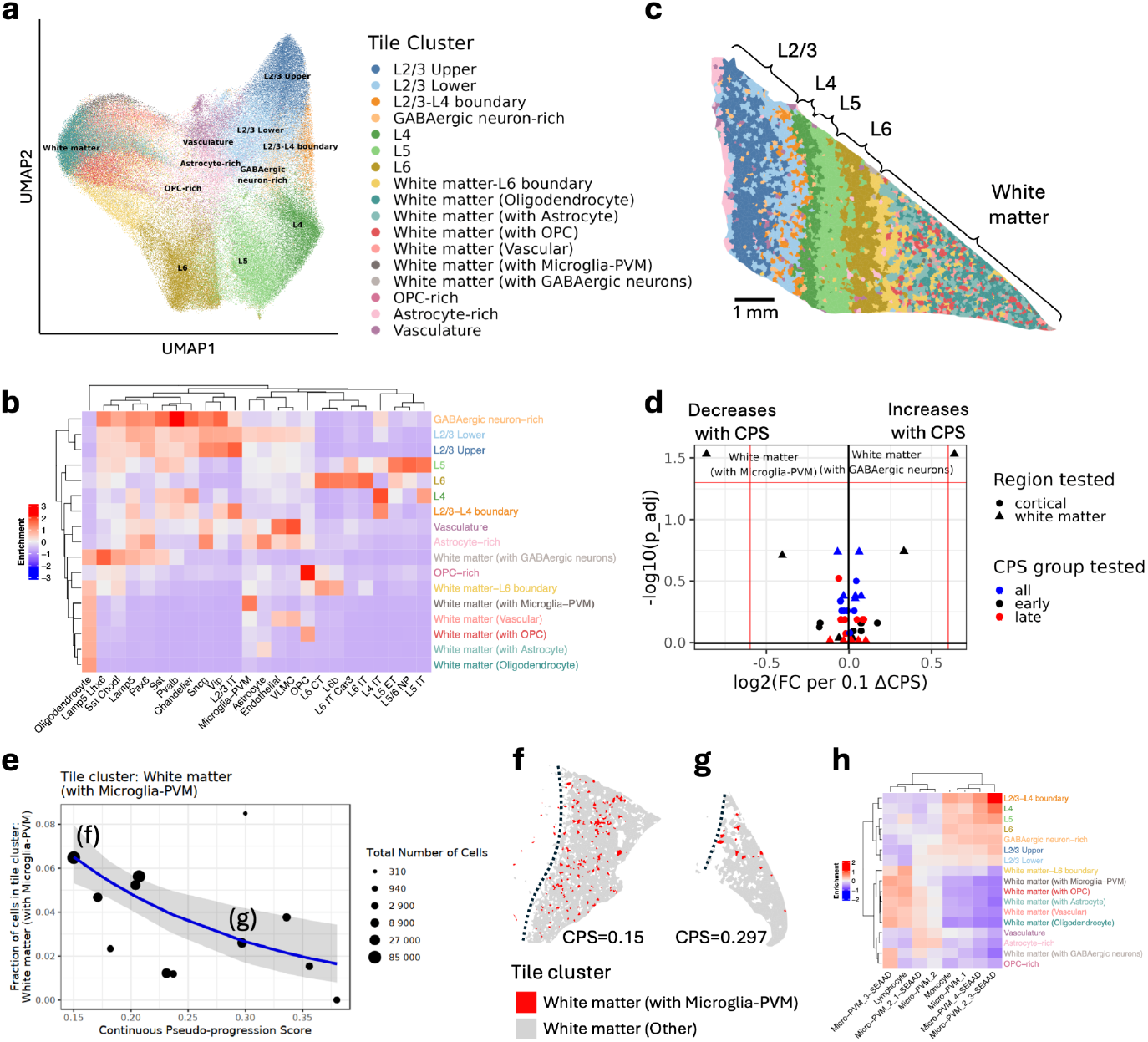
Disease-associated niches in early Alzheimer’s disease progression. **a**, UMAP visualization of Tessera tiles (each point represents a tile), with clusters identified by Leiden clustering and named according to their cell type composition. **b**, Enrichment of cell types within each Tessera tile cluster. Red (blue) indicates a cell type is found within a tile cluster more (respectively, less) often than expected by chance if cells were uniformly distributed. **c**, Spatial localization of tile clusters within a representative sample. **d**, Volcano plot of association tests between tile cluster abundance as a fraction of the cortex or white matter and increasing disease progression (Continuous Pseudo-progression Score, CPS). Association tests were conducted using a quasi-poisson GLM model. Tests were run using all donors, only using early donors (CPS ≤ 0.4), or only using late donors (CPS > 0.4). P-values were FDR-corrected using Benjamini-Hochberg correction. **e**, Association between the fraction of white matter cells in the microglia-PVM-enriched white matter tiles and CPS for early donors (CPS ≤ 0.4). Each point represents samples from a single donor, where the point size corresponds to the total number of white matter cells from that donor. **f**,**g**, Two representative white matter samples from early donors, where microglia-PVM-enriched white matter tiles are highlighted in red. The dotted lines indicate the L6-white matter boundary. **h**, Enrichment of microglia-PVM supertypes within each Tessera tile cluster. Red (blue) indicates a supertype is found within a tile cluster more (respectively, less) often than expected by chance if all microglia-PVM cells were uniformly distributed across tile clusters.

### Germinal center heterogeneity in reactive lymph node

We next asked whether Tessera could accurately isolate structures in tissues with more dynamic organization. We chose the lymph node, as it presents two challenges. Firstly, immune cells in a lymph node organize into densely packed aggregates, making it challenging to define discriminative boundaries. Secondly, germinal centers, the active sites of antigen presentation and B cell selection in the lymph node, form and dissipate in an asynchronous temporal process in response to an immunological challenge (e.g. infection). Thus, we *a priori* do not know the location, number, or maturity of germinal centers in one lymph node. Our aim is to identify germinal centers at multiple stages of maturity and characterize their cellular heterogeneity.

We applied Tessera to one reactive human axillary lymph node profiled with a 4,624-gene Xenium panel. We first identified 708K high quality cells (median 255 transcripts/cell) and grouped them into 40 major cell types (**Fig S8a**) and immune subtypes (**Fig S8b,c**) using integrative label transfer (**Methods**) with a labeled scRNAseq lymph node dataset^17–20^. Tessera grouped the cells into 23K tiles (median 34 cells/tile, **Fig S8d**). Clustering Tessera tiles, we identified 24 distinct region types (**Fig 6a, Fig S8e,f**), which fall into four classes (**Fig S8g**): lymphoid follicles (**Fig 6b**) with germinal center B cells and follicular dendritic cells (FDCs), T cell cortex (**Fig 6c**) with CD4^+^ and CD8^+^ T cells and migratory CCR7^+^ dendritic cells, medullary cords (**Fig 6d**) of lymphatic vessels and plasma cells, and other regions (**Fig 6e**) in the interstitium and lymph node capsule.

**Figure 6.**
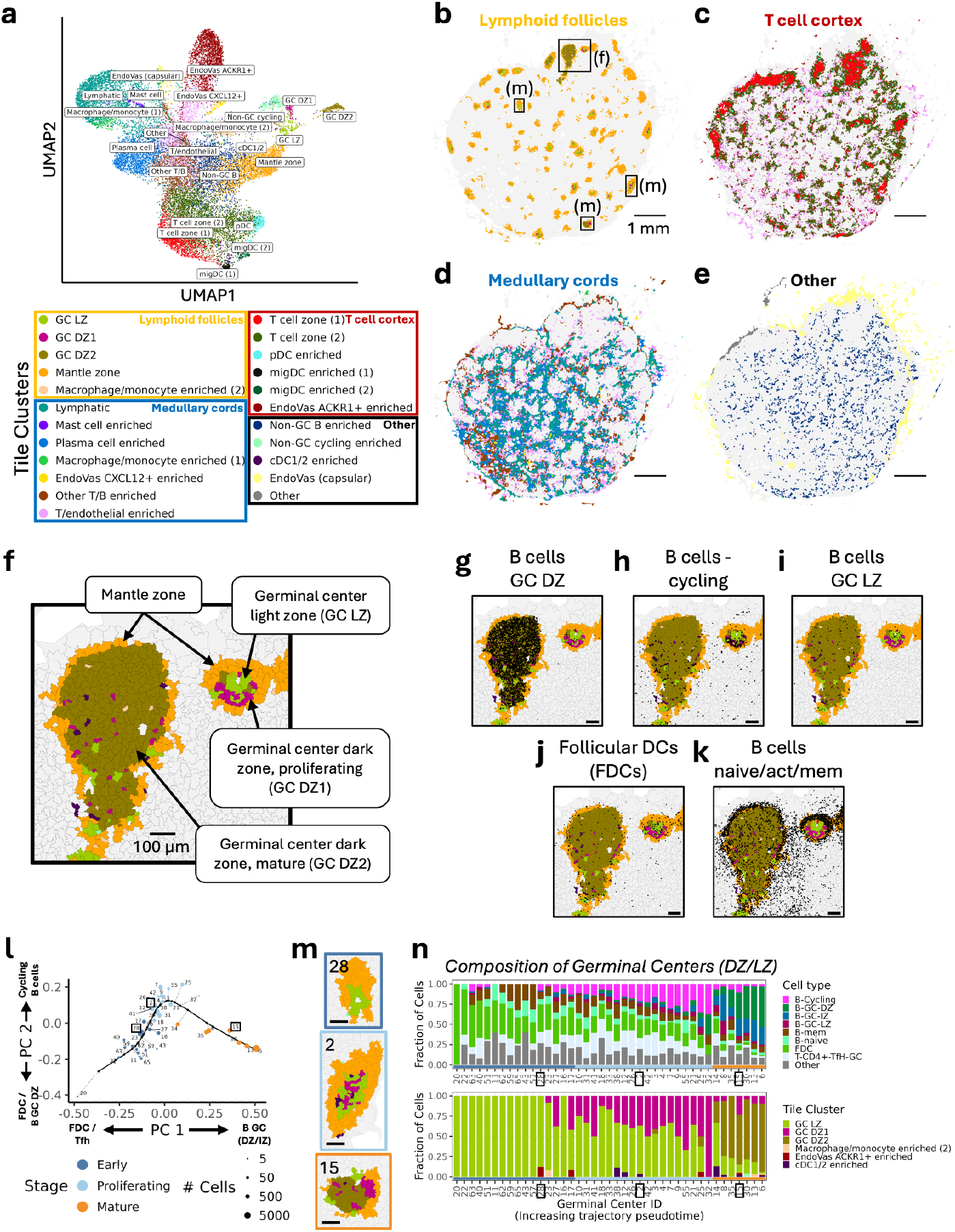
Germinal center heterogeneity in a reactive human lymph node. **a**, UMAP visualization of Tessera tiles (each point represents a tile), with clusters identified by Leiden clustering and named according to their cell type composition (**Fig S8f**). Tile clusters form distinct groups based on colocalization (**Fig S8g**), including lymphoid follicles (yellow), T cell cortex (red), and lymphatic tissue (blue). **b-e**, Spatial localization of tile clusters, split into the four groups indicated in (**a**). **f**, Tessera tile clusters for the inset in (**b**), containing multiple germinal centers. Only the mantle zone and three germinal center tile clusters are colored, with all other tile clusters shown in grey. **g-k**, For the same region as (**f**), the locations of specific subsets of cells (black points) are shown (scale bar = 100µm). **l**, Principal curve based on the cell type composition of the germinal center (light/dark zone tiles) for 41 individual germinal centers (defined in **Fig S10a-c**). PCs were computed from the cell type composition of each germinal center core, with 10 dimensions used for computing the principle curve. **m**, Spatial visualization of the tile clusters for three different germinal centers, representative of early, proliferating, and mature stages from the trajectory in (**h**). The colors match the tile clusters in (**a**). **n**, Cell type and tile cluster composition for all 41 germinal centers (light/dark zone regions), ordered according to the trajectory in (**l**).

Within the lymphoid follicles, Tessera regions reflect known substructures (**Fig 6f**): germinal center dark and light zones, enriched in germinal center B cells, cycling B cells and follicular dendritic cells (**Fig 6g-j**), and the peripheral mantle zone, a reservoir of naive and memory B cells (**Fig 6k**). Within the germinal center (GC), we found spatially separated light and dark zones. GC light zones, the site of antigen presentation, were enriched in CD83^+^ light zone B cells, CR2^+^CXCL13^+^ antigen-presenting FDCs, and CD4^+^ T follicular helper (Tfh) cells (**Fig 6i,j, S8e, S9a,d**,**g**). GC dark zones, the site of B cell mutation and selection, were divided into DZ1 and DZ2. Both contained DZ-specific *SERPIN9A* and *BACH2* expressing B cells, while DZ1 was specifically enriched in *MKI67*^+^ proliferating B cells (**Fig 6g,h, S8e, S9a-f,h,i**). We reasoned that DZ1 represents an active area within the dark zone, in which B cells were positively selected and given signals to proliferate. We expected each germinal center to contain both light and dark zones, but notably, we observed different mixtures of LZ and DZ regions across individual GCs (**Fig 6b,f**).

We therefore sought to define the heterogeneity of germinal centers in terms of their LZ, DZ, and cellular compositions. We first collapsed contiguous GC tiles into 41 distinct germinal centers (median 176 cells per center), each surrounded by a mantle zone (**Fig S10a,b**). We next used pseudotime trajectory analysis (**Methods**) to place individual germinal centers on a continuum (**Fig 6l, S10c**). Germinal centers on the “early” side of the continuum contained mostly LZ regions and FDCs (33% of cells in early GCs vs 20% and 4% in proliferating and mature), with few B cells (33% of cells in early GCs vs 48% and 82% in proliferating and mature) (**Fig 6m,n**). In the middle of the continuum, germinal centers gained more cycling B cells, potentially reflecting the highest level of B cell selection activity (**Fig 6m,n**). At the “late” side, germinal centers acquired more DZ regions and *BACH2*^+^ GC B cells (61% of cells in mature GCs vs 2% and 6% in early and proliferating, **Fig 6m,n**). Based on the progression from FDC-enriched through proliferative B-cell centers, we hypothesized that this continuum reflects germinal center maturity. Consistent with this hypothesis, the physical size of follicles also increased along the continuum, from 50 cells (median, 32-159 IQR) in “early” to 1,033 cells (median, 597-1,243 IQR) in mature follicles (**Fig S10d**). In contrast, neither the cell composition nor the physical size (median 821 cells, 579-1,247 IQR) of mantle zones varied from follicle to follicle along the continuum (**Fig S10d-f**). Together, these results suggest that we may be able to estimate the maturity of lymph node follicles by accurately segmenting the germinal center from the mantle zone and quantifying GC cellular composition.

### Anti-tumor immunity hubs in lung cancer

We next used Tessera tiles to stratify patients from a large heterogeneous cohort into clinically meaningful subgroups. We chose a dataset that used 35-plex imaging mass cytometry (**Fig 7a**), a spatial proteomics platform, to profile lung cancer biopsies from a cohort of 416 patients with a wide range of overall survival outcomes^21^. Our approach was to define shared spatial structures across all patients and cluster patients into multiple groups based on biologically important structures (**Fig 7b**). We expected these groups to represent novel molecular subtypes with distinct survival patterns.

**Figure 7.**
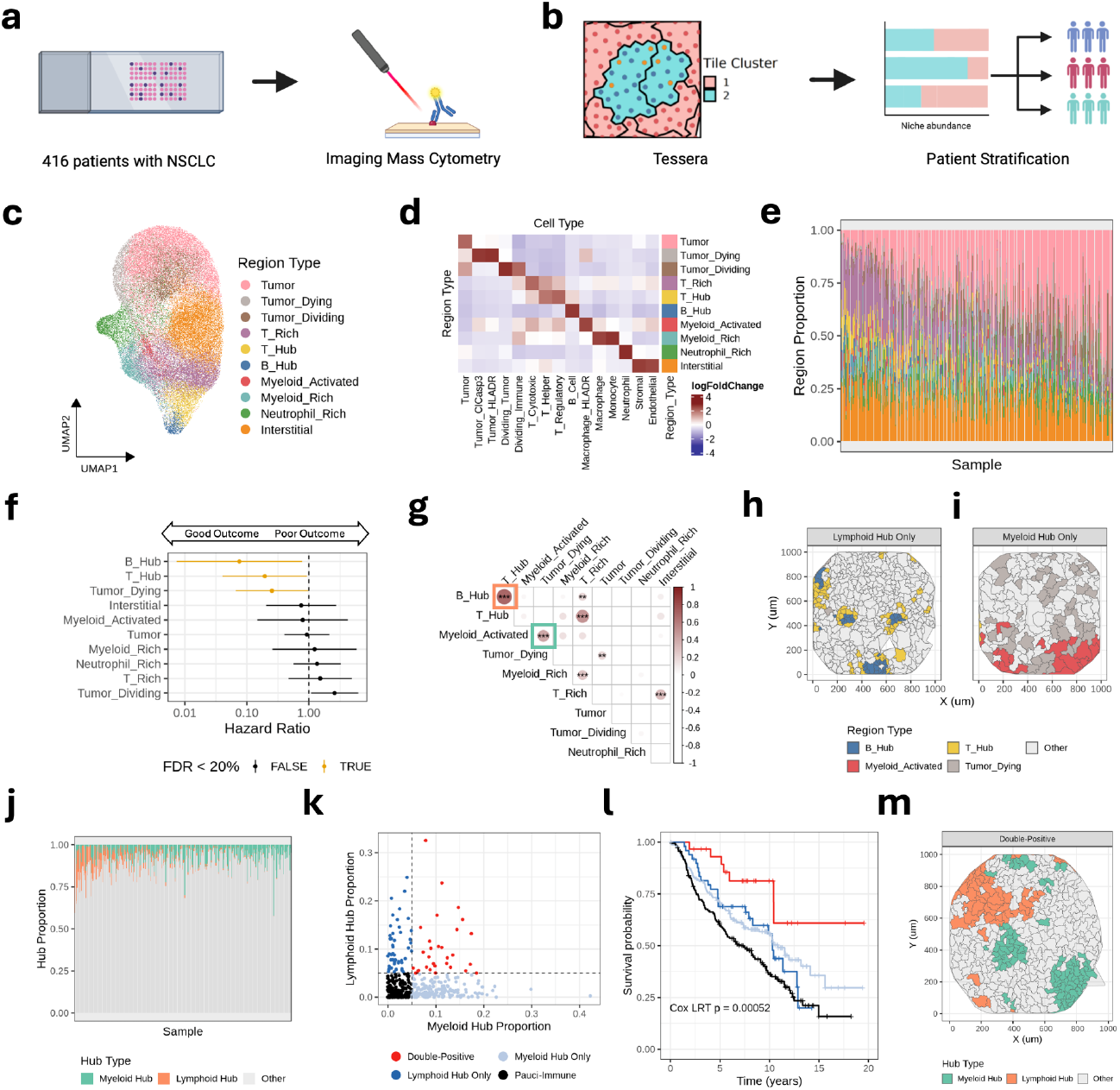
Anti-tumor immunity hubs in lung cancer. Data-driven stratification of lung cancer patients with immune-enriched Tessera tile regions. **a**, We analyzed a cohort of 416 patients with non small cell lung cancer (NSCLC), profiled with imaging mass cytometry spatial proteomics. **b**, Our analytical approach is to define distinct regions of tumor microenvironment and use abundance of these regions to stratify patients. **c**, Integrative UMAP of Tessera tiles, labeled by tile region types. **d**, Cell type compositions of tile region types. **e**, Proportions of region types across 385 high-quality samples. **f**, Forest plot shows association of sample-region abundances with patients’ survival outcomes. Hazard ratio and one-tailed p-value determined by multivariate Cox proportional hazards regression and FDR by Benjamini Hochberg. Lines represent 95% confidence intervals after adjusting for sex, age, smoking status, BMI, and disease stage. Color denotes significant (FDR<20%) association with positive outcome. **g**, Pearson correlation matrix of normalized region abundance. Stars denote significance: ***p<10^−3^, **p<10^−2^. **h**,**i**, Representative examples of samples with lymphoid (**h**) and myeloid (**i**) anti-tumor hubs, highlighting the four region types that define these hubs. **j**, Per sample proportions of anti-tumor hub types. **k**, Definition of four cohort groups based on the per sample proportion of myeloid (x-axis) and lymphoid (y-axis) anti-tumor hubs. **l**, Corresponding Kaplan-Meier survival curve for the four groups. P-value based on likelihood ratio test for 4-group term in multivariate Cox regression. **m**, Representative spatial plot of sample in the double-positive subgroup, highlighting the colocalization of lymphoid and myeloid anti-tumor hubs.

Before applying Tessera, we first identified high quality cells across all samples. Using QC and cell typing based on integrative clustering (**Methods**), we labeled 1,493,455 high-quality cells (**Fig S11a**) across 385 high-quality samples (**Fig S11b**) (**Methods**) with 8 cell lineage labels (**Fig S11c**) : MPO^+^ neutrophils, aSMA^+^ stromal cells, CD31^+^ endothelial cells, CD20^+^ B cells, CD3^+^ T cells, CD68^+^ macrophages, CD14/CD16^+^ monocytes, and PanCK^+^ tumor epithelium. Using functional markers, we then subtyped T cells into CD8a^+^ cytotoxic T cells, FOXP3^+^ Treg, and CD4^+^ T helper cells, macrophages into HLA-DR^hi^ and HLA-DR^lo^ states, and tumor cells into cleaved Caspase 3^+^ apoptotic, HLA-DR^+^ activated, Ki67^+^ dividing, and TTF1^+^ general tumor subtypes (**Fig S11c**). With Tessera, we divided these cells into 101,288 spatial tiles, with a mean of 14.7 cells per tile (**Fig S11d**). We clustered and labeled the tile clusters into 10 region types (**Fig 7c**), with distinct cell type compositions (**Fig 7d**). Three regions were primarily enriched with tumor cells (Tumor, Tumor_Dying, Tumor_Dividing), six with immune cells (T_Rich, T_Hub, B_Hub, Myeloid_Activated, Myeloid_Rich, Neutrophil_Rich), and one with stromal and endothelial cells (Interstitial). These ten regions are present broadly across all 385 samples (**Fig 7e**) and thus serve as a plausible basis for patient-to-patient comparison. We next asked which regions may be associated with clinical outcome. We performed survival regression analysis, accounting for sex, age, smoking status, BMI, and disease stage as covariates (**Methods**), and found three regions significantly (FDR<20%) more abundant in patients with good outcomes: B_Hub, T_Hub, and Tumor_Dying (**Fig. 7f**). We next asked whether these regions are spatially distinct or represent components of larger regions. Using abundance covariance analysis (**Fig 7g**), we found that patients with many B_Hub tiles also had more T_Hub tiles (**Fig S11e**, spearman r=0.68, p<2.2×10^−16^) and the two regions colocalized in space (**Fig 7h, S11g**). Similarly, the Tumor_Dying niche covaried (spearman r=0.40, p=2.5×10^−16^, **Fig 6g, S11f**) with the Myeloid_Activated niche and colocalized in space (**Fig 7i, S11h**). Given these co-abundance and colocalization patterns, we reasoned that these four regions formed two spatially distinct anti-tumor hubs: one lymphoid anti-tumor hub, with spatially separated B cell and T cell compartments and one myeloid anti-tumor hub, with actively dying tumor cells expressing apoptotic marker cleaved Caspase 3 abutting activated HLA-DR^+^ macrophages. The abundance of these two anti-tumor hubs varied across the cohort (**Fig 7j**) and thus may explain different survival outcome patterns in patients.

We reasoned that the presence of lymphoid and myeloid hubs may define distinct molecular subtypes of lung cancer. Based on the normalized abundance (**Methods**) of each hub type, we grouped patients into four groups (**Fig 7k**): pauci-immune (n=157) with neither, lymphoid hub only (n=49), myeloid hub only (n=149), and double-positive (n=30). These patient groups exhibited significant differences in overall survival (**Fig 7l**, p=0.00052). The pauci-immune group had the worst outcome (median survival = 7.4 yrs), while patients with exactly one of the anti-tumor hubs fared better (p=0.026) (**Fig S11i**). However, we found no significant difference (p=0.38) between the lymphoid hub only (median survival = 10.3 yrs) and myeloid hub only (median survival = 10.9 yrs) groups (**Fig S11j**). Patients with both niches had the best survival outcomes (median survival >20 yrs), better than both the lymphoid hub only (p=0.0027) (**Fig S11k**) or myeloid hub only (p=0.0085) groups (**Fig S11l**). Importantly, every spatial sample in this study captures a small region of ∼1mm^2^. Therefore, the presence of both lymphoid and myeloid hubs in one sample suggests the colocalization between the two niches (**Fig 7m, S11m**). We posit that the “super-responder” group represents patients with a coordinated, two-pronged anti-tumor immune response.

## Discussion

Spatial transcriptomics in human tissue promises a new wave of insights into human disease by defining how cells communicate within tightly controlled tissue niches. Technological advances have focused on spatial resolution^22^, allowing us to profile spatial gene expression at 100µm, then 50µm, and now (subcellular) 0.2µm. Algorithms to analyze data from these platforms have also emerged to leverage this increased resolution. CellCharter^4^ and STAGATE^6^ exemplify computational methods to automatically define tissue domains of different sizes and shapes. Unfortunately, the same tools these algorithms use to define spatially contiguous structures also make them prone to blurring true edges between adjacent structures. Tessera is the first algorithm to explicitly model the natural boundaries of tissue structures and incorporate these boundaries into spatial analysis. Tessera groups cells into spatially contiguous, boundary-respecting tiles that can be clustered for downstream detection of biologically meaningful tissue domains. Tessera tiles represent a conceptually novel way to represent cells in space. By design, these tiles represent the lowest level of multicellular spatial organization. While we focus on clustering tiles for domain detection, these tiles are amenable to a number of downstream spatial analyses, such as non-negative matrix factorization to discover spatially patterned gene programs, registration of serial sections with tile-based anchors, and spatially constrained ligand-receptor analysis to infer accurate cell-cell communication networks. Finally, Tessera tiles provide a robust basis for comparison of multiple samples. By running Tessera on harmonized cell embeddings, we recover biologically comparable tissue regions across donors and find disease-associated regions by comparing case vs control donors.

The quality of Tessera tiles depends on several factors. First, Tessera estimates spatial gradients around cells and thus depends on reasonable cell segmentation. While Tessera is robust to errors in cell boundary estimation, over-segmentation (i.e. splitting one cell into many) will misspecify the extent of spatial gradients and lead to incorrect tile boundaries. The second factor is which cell features are passed to Tessera. For instance, with too few principal components, Tessera will find boundaries between regions with different cell types but miss boundaries related to changes in cell activation states. The third factor is quality control. We found that when Tessera tiles are consistently small in one region, this often reflects a low quality region (e.g. tissue fold) whose noisy gene measurements result in inconsistent gradients and small tiles.

Tessera represents a novel paradigm to perform segmentation in spatial omics data. At its core, Tessera inputs a set of points in space (i.e. cells) and divides them into molecularly interpretable groups of points (i.e. tiles). Thus, Tessera can be applied to spatial data at any resolution, such as subcellular pixels measured by spatial sequencing technologies (i.e. Visium HD^23^ and Stereo-Seq^24^) and even individual transcripts in space, to define subcellular compartments and perform more robust cell segmentation. Importantly, we built Tessera on a foundation of robust and scalable computational tools, including anisotropic bilateral filtering, with a rich history in computer vision^25^, and Discrete Morse Theory^9,10^, an emerging tool from topological data analysis, a broad computational field with multiple successful tools imported into genomics (e.g. UMAP^26^, persistent homology^27,28^). Importantly, these foundational tools readily scale to big datasets (i.e. billions of points) and can be adapted to more complex analysis scenarios, such as spatial multi-omics and 3D imaging. We hope that Tessera serves both as a useful tool for automated tissue domain detection and as a broader paradigm for downstream spatial analysis and methods development.

## Methods

### 1. Tessera algorithm

The Tessera algorithm takes as input single cells (or pixels) with spatial coordinates **x**_*i*_=(*x*_*i*_, *y*_*i*_) ∈ ℝ ^2^and cell embeddings **z**_*i*_ ∈ ℝ ^*d*^(or transcript counts) for each cell *i*. The output is a segmentation of adjacent cells into tiles with a user-controlled size parameter. Boundaries between tiles align with where cell composition and gene expression change the most within the tissue. Segmentation using the Tessera algorithm has four main steps^29^, detailed in the later sections:

#### 1. Constructing inputs

A triangle mesh is constructed using Delauney triangulation and pruned to eliminate long edges. If transcript counts are provided for each cell instead of embeddings, then cell embeddings are computed using principal component analysis (PCA).

#### 2. Gradient estimation

Gradients are calculated at each vertex by considering the difference in cell embeddings between each cell and its neighbors in the mesh. These gradients are smoothed using anisotropic bilateral filtering, and then gradients are defined for edges and triangles in the mesh by averaging the vertices that each edge or triangle contains.

#### 3. Tissue segmentation using discrete Morse theory (DMT)

A scalar field is defined by taking the magnitude of the total gradient at each vertex, edge, and triangle. Then DMT-based segmentation is performed by constructing a maximum spanning forest on the triangles and a minimum spanning forest on the vertices^9^. Separatrices that partition cells into tiles of homogeneous composition are defined by tracing paths between critical points, specifically between saddle edges and maximum triangles.

#### 4. Hierarchical agglomeration

Tiles from DMT-based segmentation are merged using single-linkage agglomerative clustering to obtain tiles containing a number of cells between a user-provided minimum and maximum value. Pairs of adjacent tiles are scored according to their transcriptional similarity, compactness of shape after merging, and number of cells, in order to prioritize favorable merges in each agglomerative clustering step.

### 1.1 Constructing inputs

If cell embeddings are not already provided, then they are computed using the transcript counts by log-normalization and principal component analysis (PCA). Specifically, total counts per cell are normalized to the median across all cells and log1p-transformed. Each gene feature is then Z-scored, with zero mean and unit variance. The top *d* principal components are computed using all genes, to obtain a cell embedding matrix **Z**∈ ℝ^*d*×*n*^, where *d* is the embedding dimension and *n* is the number of cells (or pixels). Alternatively, cell embeddings **Z**∈ ℝ^*d*×*n*^, can be provided directly by the user. In this case, cell embeddings can be computed in a number of ways, including different approaches to pre-processing and linear or non-linear dimensionality reduction. A triangle mesh is computed from the cell (or pixel) locations using Delauney triangulation, where each cell is a vertex in the mesh. Each edge in the mesh connects two adjacent vertices and two adjacent triangular faces. Spurious connections between distant cells within the tissue are pruned by removing edges that are longer than a given distance. Cells that become disconnected from the rest of the sample after pruning are not assigned to any tiles during the segmentation process.

The mesh can be viewed from the perspective of vertices connected by edges—this forms the primal graph of vertices and edges. The mesh can also be viewed from the perspective of triangles joined by shared edges—this forms the dual graph of triangles and edges. Each edge participates in both the primal graph and the dual graph^29^.

### 1.2 Gradient estimation and smoothing

First, we compute a spatial gradient field **F**_*i*_∈ ℝ^2×*d*^, at each vertex (cell) *i*:

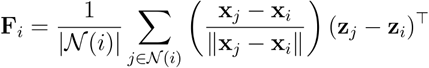

The gradient field is the average gradient in expression of each embedding dimension between the index cell *i* and its neighbors *j* ∈ 𝒩(*i*). The distance between neighboring cells is normalized to be unit distance, so that only the direction from each cell to its neighbors matters.

The gradient field at each vertex is smoothed using bilateral filtering^30^, in which the smoothed gradient is computed as the weighted average of the neighbors’ gradients, considering both spatial distance and also similarity in gradient values:

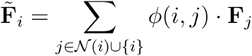

Bilateral filtering is an effective method for reducing noise while preserving discontinuities^30^. The (unnormalized) weight 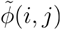 of each neighbor is computed as the product of two scores:

#### 1. Distance score 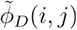

Generally, more distant neighbors have less weight. There are two methods for calculating the distance between adjacent cells:

a. If the “Euclidean” method is used, then the distance between adjacent cells is computed as 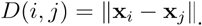
b. If the “projected” method is used (default), then the distance between adjacent cells is computed as 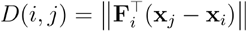. This is an anisotropic filter that accounts for the expected change in gene expression towards the direction of the neighboring cell. Using the gradient field, it is calculated as the total derivative in the direction of the neighbor.

We then compute a Gaussian transformation of the spatial distance of cell from its neighbor *j*, so that more distant neighbors have less weight. For both distance methods, the relative cell weight is defined as

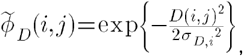

where 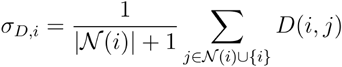 is the average distance.

#### 2. Similarity score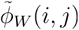

Generally, neighbors with more similar gradients have greater weight. There are two methods for calculating similarity:

a. If the “Euclidean” method is used, then the dissimilarity between adjacent cell gradients is computed as the Frobenius norm 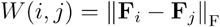.
b. If the “projected” method is used (default), then the dissimilarity between adjacent cell gradients is computed as the cosine distance between a cell’s gradient field and its neighbor’s gradient field: 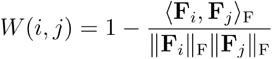, where ⟨ ·, · ⟩_F_ is the Frobenius inner product. We then compute a Gaussian transformation of the gradient dissimilarity, so that more dissimilar neighbors have less weight. The relative cell weight is defined as

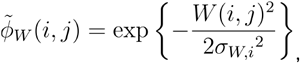

where 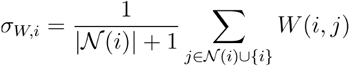 is the average dissimilarity.

Finally, the normalized weight for each neighbor is

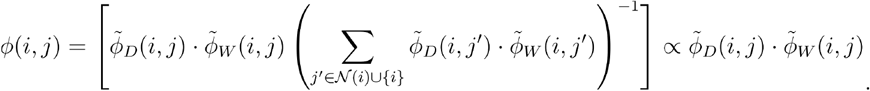

The number of bilateral filtering iterations can be selected by the user (“smooth_iter” parameter). Typically, a small number of bilateral filtering steps (between 1 and 5) is sufficient to smooth gradients effectively.

After smoothed gradient fields 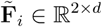 have been computed at each vertex, gradients are computed for each triangle as the average of its vertices’ gradients. Each edge in the mesh connects two adjacent vertices (an edge in the primal graph of vertices and edges) and two adjacent triangular faces (an edge in the dual graph of triangles and edges). The primal and dual edge gradients are computed as the sum of the gradients for the two points and two triangles, respectively, that are associated with each edge.

### 1.3 Tissue segmentation using discrete Morse theory

For each vertex, primal/dual edge, and triangle, scalar field values *f* are computed as the Frobenius norm of the spatial gradient field 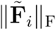 (**Fig S2a**). High values of *f* indicate regions where the cell composition or gene expression is changing most sharply, and we seek to partition cells (vertices) into tiles whose boundaries align with edges that have high values of *f*. To do this, we apply an algorithm based on discrete Morse theory that efficiently identifies salient edges with maximal curvature, called topological separatrices, by constructing the Morse-Smale complex^9,29,31^. The term “curvature” derives from previous applications to 3D meshes, which aim to draw boundaries that track with the curvature of the surface of an object in 3D space. In our setting, curvature is defined in transcriptional space and denotes the total change in gene expression in a local spatial neighborhood.

Specifically, we construct a directed minimum spanning forest on the primal graph of vertices and edges using Kruskal’s algorithm (**Fig S2b-c**). Edge weights are given by *f*. Critical vertices (local minima; or, more precisely, an endpoint of an edge that is a local minimum) are used as roots for each tree in the forest, and edges that would bridge two trees with different critical vertex roots are marked as possible saddle edges.

Similarly, we construct a directed maximum spanning forest on the dual graph of triangles and edges using Kruskal’s algorithm (**Fig S2d-e**). Edge weights are given by *f*. Critical triangles (local maxima; or, more precisely, a triangle that contains an edge that is a local maximum) are used as roots for each tree in the forest, and edges that would bridge two trees with different critical triangle roots are marked as possible saddle edges.

To construct separatrices from the primal and dual spanning forests, we trace all paths from saddles to dual critical triangles (which are local maxima in the value of *f*). Saddle edges are the edges that are possible saddles in both the primal and dual spanning forests (**Fig S2f-g**). Because the dual spanning forest is a maximum spanning forest on triangles in the mesh (**Fig S2h**), the paths that are traced between saddles and critical triangles (maxima) will follow ridges with high values of *f* (**Fig S2i-k**), separating points into components (tiles) with the strongest boundaries (**Fig S2l**).

The separatrices partition the cells (vertices) into distinct connected components, which we call “tiles”. The number of cells assigned to each tile is determined by the number of local minima in the value of *f* (**Fig S2j**), which depends on the amount of smoothing that is performed. In practice, we often observe that the number of cells per tile is between 1 and 20. Each tile 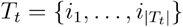 is represented by an embedding vector

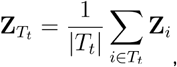

which is the average of the tile’s cell embeddings.

### 1.4 Hierarchical agglomeration

Because the initial tiles obtained from DMT-based segmentation are often small due to fluctuations in the scalar field values, we agglomerate adjacent tiles into larger tiles until we obtain tiles within a user-specified range of sizes (by default, between 5 and 50 cells). Pairs of adjacent tiles are scored according to their transcriptional similarity, compactness of shape after merging, and the number of cells in order to prioritize favorable merges in each agglomerative clustering step. The score for each pair of adjacent tiles is computed as the product of the following three factors, where higher scores favor merging:

#### 1. Transcriptional similarity

We compute the dissimilarity between adjacent tiles *T*_*s*_ and *T*_*t*_ as the Euclidean distance *d*(*T*_*s*_, *T*_*t*_)= ‖**Z**_*s*_ − **Z**_*t*_‖. To obtain a null distribution for the dissimilarity between tiles with similar gene expression, we fit a 2-cluster Gaussian mixture model (GMM) to the values for *d*(*T*_*s*_, *T*_*t*_)across all adjacent tiles *T*_*s*_ and *T*_*t*_. The cluster with smaller mean (mean *μ* and standard deviation *σ*) defines the null distribution. Finally, we calculate the transcriptional similarity as

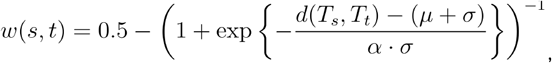

which ranges from − **0.5** to 0.5. If adjacent tiles are very dissimilar (*d*(*T*_*s*_, *T*_*t*_) ≫ *μ* + *σ*), then *w*(*s,t*) ≈ − **0.5**. If adjacent tiles are similar (*d*(*T*_*s*_, *T*_*t*_) ≫ *μ* + *σ*), then *w*(*s,t*)>0. By default, tiles are merged as long as and the number of cells in each tile is not too large. The value of (default) controls the strength of merging based on transcriptional similarity, where higher values around 2 correspond to more liberal merging, and lower values around 0.2 correspond to more conservative merging.

#### 2. Size S score for number of cells in the merged tile

For adjacent tiles *T*_*s*_ and *T*_*t*_, we define the size score

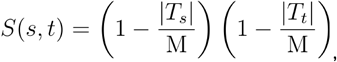

where M is a user-defined parameter for the maximum tile size. The score *S*(*s,t*)ranges from 0 to1. If merging two tiles would have a total number of cells ≥M, then *S*(*s,t*)= − ∞, which prevents merging.

#### 3. Compactness score ΔC

Shape compactness is defined as the ratio between area and perimeter-squared: 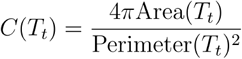, where a circle has the maximum possible compactness of 1. We define the compactness score associated with merging two adjacent tiles *T*_*s*_ and *T*_*t*_, to be

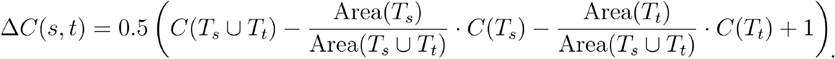

This score ranges between 0 and 1.

We perform single-linkage agglomerative clustering to aggregate tiles into larger tiles with up to M cells (where M is user-defined). Every pair of adjacent tiles *T*_*s*_ and *T*_*t*_ has an associated merging score *W*(*s,t*)=*w*(*s,t*) ·*S*(*s,t*) · Δ*C*, where higher scores favor merging. Merging is conducted greedily, where we perform the merge with the highest score at each step, and then update the scores for the newly formed tile. Merging stops when *W*(*s,t*)<0for all adjacent tiles, which happens when neighboring tiles have very dissimilar gene expression *w*(*s,t*)<0or when tiles reach the size limit M.

Finally, to ensure that all tiles have at least m cells (where m is also user-defined), we perform one more round of agglomerative clustering, considering only pairs of adjacent tiles where at least one tile has < *m* cells. Additionally, we use a transcriptional similarity score of

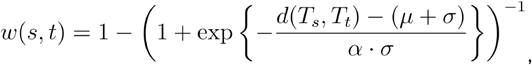

which ranges from to, instead −0.5of to 0.5, to allow even more dissimilar tiles to merge if the minimum size threshold is not yet reached. (Note that as a result of this final round of clustering, it may be possible to obtain some tiles containing more than M cells, because very small tiles were merged into a larger one.)

The final result is a partition of the original collection of cells into *i* ∈{1,…,*n*}tiles 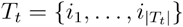, each with associated embedding vector

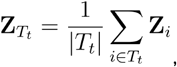

which is the average of the tile’s cell embeddings. Additionally, if transcript counts are provided for the input cells, then each tile also has associated transcript counts, representing a local pseudo-bulk of the single cells in the spatial transcriptomics data. This is technically similar to pseudobulking counts in the meta-cell approach^32^. However, while the goal of meta cells is to collapse counts for groups of transcriptionally similar cells, Tessera tiles often contain cells of different types and so represent heterogeneous mixtures of distinct cell programs.

### 2. Multi-sample analysis

In order to integrate spatial transcriptomics data from multiple samples and accurately find shared cell types or spatial domains, we must account for batch effects, as in other single-cell data analysis^33^. The Tessera pipeline readily accommodates existing single-cell integration methods^34^.

First, before running Tessera, we apply Harmony v1.2 to remove cell type-specific batch effects from the cell inputs^35^. Harmony v1.2 scales readily to 10s of millions of cells and hundreds of batches. It produces harmonized PC embeddings **Z**_*i*_ for each cell *i*, which we use as inputs for Tessera. As a result, the tile embeddings 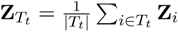 are computed using cell embeddings that are already well-integrated and no longer confounded by batch effects. These tile embeddings can be used for downstream tasks such as joint clustering, without any need for further batch correction. We stress the importance of only applying Harmony to cells and not directly to tiles. While we can technically treat tiles as individual units of observation and apply Harmony to the PCA embeddings of tiles, the interpretation of Harmony corrected embeddings is often wrong. Briefly, Harmony correction of tiles cannot distinguish between batch-specific noise applied to cell types and batch-specific differences in tile composition. We only want to correct for the first and not the second. As a result, Harmony applied to tiles merges tiles across batches with slightly different cell compositions. In practice, this often breaks the spatial coherence of tiles in space and makes it difficult to robustly define larger tissue regions in space. Alternatively, other integration methods for multi-modal single-cell data could also be used to obtain integrated cell embeddings that are used as inputs for Tessera^34,36^.

### 3 Spatial domain analysis

#### 3.1 Clustering Tessera tiles

Once Tessera tiles are constructed, they can be clustered using their tile embeddings just as in single-cell clustering analysis. While many different strategies are possible, we either 1) used Seurat’s FindNeighbors and FindClusters functions to construct a tile nearest neighbor graph and cluster using the Louvain modularity clustering algorithm, or 2) constructed a fuzzy nearest neighbor graph using the UMAP algorithm and then clustered using the Leiden algorithm (as done in the scanpy framework)^37,38^. Just as in clustering and refinement of cell types in scRNA-seq data, tile clusters can be further refined by sub-clustering, in which a subset of tiles are selected and clustered on their own^39^. In some instances, it can be helpful to recompute cell and tile embeddings such as principal components (PCs) on a selected subset of cells and tiles before subclustering in order to enrich for variability within that subset of the data.

#### 3.2 Cell composition of Tessera tile clusters

Once we identified tile clusters, we computed the enrichment of cell type *i* in tile cluster *j* as 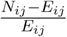, where *N*_*ij*_ is the observed number of cells of cell type *i* within tile cluster *j*, and 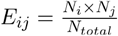 is the expected number if cells were randomly distributed across tile clusters (*N*_*i*_ is the number of cells of cell type _*i*_, *N*_*j*_ is the number of cells assigned to tile cluster *j*, and *N* _*total*_ is the total number of cells). In the tile cluster-cell type enrichment heatmaps, these values are sgn (*x*) ·log(1+ | *x* |-transformed for visualization purposes.

#### 3.3 Colocalization of Tessera tile clusters

Colocalization of tile clusters was computed by counting the number of times a tile belonging to cluster *j*_1_ was adjacent to a tile of cluster *j*_2_ across the entire dataset. We then simulated a null distribution by permuting the cluster label for all tiles and recomputing the number of *j*_1_-*j*_2_ neighbors. This null distribution modeled the case in which tile clusters were distributed randomly, controlled for the abundance of each tile cluster. Finally, we used the null distribution to compute a mean and a standard deviation to Z-score the actual number of *j*_1_-*j*_2_ neighbors. We then computed this Z-score for all combinations of *j*_1_ and *j*_2_ and plotted the results in a heatmap.

## 4. Whole mouse brain MERFISH dataset

### 4.1 Data preprocessing

We downloaded MERFISH spatial transcriptomics data from 147 coronal sections of a single adult mouse brain (animal 1) as a Seurat object from the CELLxGENE data portal (https://datasets.cellxgene.cziscience.com/c3e7611c-e5ba-4cb3-ba53-bb2fc75d74be.rds)^11^. The MERFISH data had already been processed into 4.2M cells that passed quality control. Each cell was provided with raw transcript counts from a 1,122-gene panel, spatial coordinates, and cell type annotation (25 broad cell types, and 338 subclasses).

### 4.2 Single-sample analysis

A single coronal section (C57BL6J-1.079), which contained 46,339 cells, was selected. For every cell in this section, transcript counts were log-normalized using the median number of transcripts per cell and Z-scored. The top 50 principal components (PCs) were computed across all the cells using all 1,122 genes. Next, we constructed Tessera tiles using 50 PCs and minimum/maximum tile sizes of 5-50 cells. During Tessera, three rounds of gradient smoothing were performed using the “projected” distance and similarity metrics. Embeddings (PCs) were aggregated for all cells in each tile.

Tiles were then clustered using graph-based Leiden clustering (where the nearest-neighbor graph was computed using the `uwot` UMAP similarity graph) in two stages: first, all tiles were clustered at a coarse level to produce 19 major clusters (using 30 nearest neighbors, Leiden resolution = 1.8); afterwards, tiles belonging to five of these clusters were selected and further subclustered on their own. For each round of subclustering, tiles for the specified tile cluster were isolated, 15 expression PCs were recomputed on that subset of tiles, and then clustering was performed on that subset alone (using 10 nearest neighbors, Leiden resolution 0.2-0.7). Tile clusters were annotated based on spatial localization and enrichment of previously annotated cell types and cell subclasses.

### 4.3 Multi-sample analysis

All 4.2M cells from 147 coronal sections were analyzed together. First, transcript counts were log-normalized using the median number of transcripts per cell and Z-scored. The top 50 principal components (PCs) were computed across all the cells using all 1,122 genes. Then harmonized PCs (hPCs) were computed using Harmony v1.2, integrating across the “brain_section_label” batch variable (treating each section as its own batch) for 10 iterations. Next, Tessera was run separately on each section using the hPC cell embeddings as input, with default parameters (min/max tile sizes of 5-50 cells). Within each tile, cell embeddings (hPCs) were averaged to produce tile embeddings that were integrated across all sections.

Tiles from all sections were clustered together using graph-based Leiden clustering: first, a fuzzy 30-nearest-neighbor graph was computed using the `uwot` UMAP similarity graph; afterwards, modularity-based Leiden clustering of this graph was done with resolution 1.0, yielding 36 major tile clusters. These tile clusters were annotated based on spatial localization of the clusters and enrichment of previously annotated cell types and cell subclasses. The tile clusters were named using the nomenclature of the Allen Mouse Brain Atlas and validated by comparing with coronal sections from the reference brain map (https://atlas.brain-map.org/atlas?atlas=1).

## 5. Benchmarking time and memory

Tessera and CellCharter pipelines were run on 2, 5, 10, 20, 50, 100, and 147 brain slices, corresponding to between 72,000 and 4.2 million cells. Benchmarking analysis was carried out on a Linux-based high performance computing (HPC) platform, using virtual environments with up to 200GB of memory. The available hardware included Intel Xeon E5 CPUs and Tesla V100 GPUs with 16GB of VRAM. The time elapsed was measured for each step of the Tessera and CellCharter pipelines, and the maximum memory usage was profiled using the Linux `/proc/pid/statm` file (resident set size). Running STAGATE (with the efficient STAGATE_pyG implementation) simultaneously on multiple brain slices failed due to GPU memory constraints.

CellCharter was run using 5 CPUs and 1 GPU, since using more cores made no difference in performance. The CellCharter pipeline was broken down into four steps: 1) feature normalization, 2) batch integration with scVI, 3) feature aggregation, and 4) clustering of cells with a Gaussian mixture model (GMM, implemented in PyTorch with GPU-acceleration). For feature normalization, transcript counts were log-normalized using the median number of transcripts per cell. As required for scVI, features were not Z-scored. Cells were integrated across all sections with scVI, using “brain_section_label” as the batch variable. Default settings were used for scVI with a max epochs heuristic and with early stopping. Consequently, the number of epochs decreased proportionally to the number of cells, so the scVI runtime was approximately constant regardless of the number of cells. CellCharter feature aggregation of the scVI cell embeddings was performed using default parameters, with Delauney triangulation (removing long links with percentile = 99) and aggregation of 0, 1, 2, and 3-hop neighbors (n_layers = 3). Finally, a single round of GMM clustering was performed using a fixed number of clusters (n_clusters = 32).

Tessera was run using 20 CPUs. The Tessera pipeline was broken down into four steps: 1) feature normalization and dimensionality reduction with PCA, 2) batch integration of cells with Harmony, 3) tile construction with Tessera, 4) graph-based Leiden clustering (UMAP similarity graph construction followed by modularity-based clustering). In steps 1, 2 and 4, all sections were analyzed together. In step 3, Tessera tiles were constructed on each section separately, allowing for parallelization across sections. For feature normalization, transcript counts were log-normalized using the median number of transcripts per cell and Z-scored. The top 50 principal components (PCs) were computed across all the cells using all 1,122 genes. Cells were integrated across all sections with Harmony v1.2, using “brain_section_label” as the batch variable. Default settings were used for Harmony, along with parallelization (ncores = 20). Tessera tiles were constructed using min/max tile sizes of 5-20 cells, one round of smoothing, and pruning of 99th-percentile edges (to match CellCharter parameters). Tile embeddings were computed as the average of the harmonized cell embeddings. Finally, the UMAP similarity graph of tile embeddings was constructed using 30 nearest-neighbors, and Leiden clustering of the tiles was performed using a fixed resolution = 1.0.

## 6. Benchmarking against other methods

### 6.1 Ground truth annotations

Ground truth annotations for cortical layer and ventricle regions were defined based on the localization of specific subclasses of neurons and ventricular cells. We chose to manually annotate the regions using the cell type labels rather than using the Allen Atlas Common Coordinate Framework (CCF) because this allowed us to define boundaries with single-cell precision. We often observed that while the CCF coordinates approximately described which regions are expected, they did not provide accurate boundaries between regions within specific sections.

For the cortical layers, neuron annotations were combined into 12 different labels corresponding to neurons that localize to distinct cortical layer regions. These neuron labels were then propagated to neighboring cells and any remaining “holes” of unassigned cells within the cortex were identified and annotated based on their neighbors. To avoid dilation of the manually annotated regions, only non-neuronal cells within the interior of the cortical region were re-labeled based on their neighbors.

For the ventricle regions, three ventricular cell types were selected: choroid plexus epithelial cells, ependymal cells, and tanycytes (a specialized ependymal cell type). In order to define coherent spatial domains based on these cell types, all other cells were given a distinct “other” label, and then each cell’s identity was reassigned based on voting using its neighbors’ labels. Voting-based label propagation was performed until convergence.

While our manually annotated regions accurately described the spatial localization of the selected region-specific cell types, there were still ambiguities that might lead to some disagreement even for a well-designed spatial domain detection method: 1) boundaries may not be perfectly delineated between adjacent cortical layers if the different cell types have some overlap in their locations, 2) for some regions, there are a few isolated cells that are not connected to the rest of the region (these cells are included to avoid erasing small structures), and 3) there may be valid tissue structure (e.g. blood vessels) within broader cortical regions that could merit further subdivision.

### 6.2 Metrics for benchmarking

When benchmarking different methods (Tessera, CellCharter, and STAGATE) in their ability to recapitulate known, manually annotated spatial domains, we measure two different characteristics: 1) accuracy in identifying spatial domains that match the manual annotations, and 2) coherence of the spatial domains.

The accuracy of spatial domains is quantified using an F1-score that requires detected domains to have both high recall (captures most of an annotated region’s cells in one domain) and high precision (avoids including cells from outside of the region). Because Tessera, CellCharter, and STAGATE each assign cells into distinct spatial domains in an unsupervised manner, the F1-score for an annotated region is computed across all spatial domains, and the domain with the highest score is chosen to match that region. Perfect alignment with the annotations would result in an F1-score of 1.

Spatial coherence is quantified in order to capture behavior where these methods cluster cells by cell type rather than by spatial context, resulting in cells whose spatial domains disagree with their neighbors. We quantify incoherence as the average fraction of neighboring cells assigned to a different spatial domain as the index cell. Ideally, spatial domains span broad swaths of tissue, resulting in an incoherence close to zero.

### 6.3 Settings for each method

Tessera, CellCharter, and STAGATE were each applied to one entire coronal section (C57BL6J-1.079, which contained 46,339 cells) to group cells into spatial domains.

Tessera was run using min/max tile sizes of 5-20 cells, one iteration of smoothing, and pruning parameters chosen to best match the graph construction in CellCharter and STAGATE (prune_thresh_quantile = 0.99 and prun_min_cells = 1). Tile clustering was done by constructing a fuzzy 15-nearest-neighbor graph with the `uwot` UMAP implementation. Tiles were then clustered using modularity-based Leiden clustering for a range of resolution values. Cells were assigned to their associated tile cluster to produce cell-level spatial domains for comparison with CellCharter and STAGATE.

CellCharter was run using scVI to construct cell embeddings and then aggregation of 0, 1, 2, and 3-hop neighbors (n_layers = 3). Clusters were then found using the CellCharter GMM implementation across a range of n_clusters parameters. For each value of n_clusters, up to 7 clustering solutions were generated, and the best one was selected (using the ClusterAutoK function).

STAGATE was run using the STAGATE_pyG implementation (https://github.com/QIFEIDKN/STAGATE_pyG) with default parameters to produce spatially-informed cell embeddings. These embeddings were then clustered using Leiden clustering across a range of resolution values.

### 6.4 Parameter search for cluster resolution

For each method, the parameter that will most affect the resulting spatial domains is the clustering resolution (Tessera and STAGATE Leiden clustering) or number of clusters (CellCharter GMM). The clustering resolution will affect the scale of the spatial domains that are detected. Biologically, multiple resolutions could produce valid results because there is often a hierarchical organization to tissue (e.g. the same method might capture the entire cortex as one unit at a lower clustering resolution, or separate individual cortical layers at a higher clustering resolution). To produce a valid comparison between the different methods that best matches the manually annotated regions, we scanned a wide range of clustering resolutions (res=0.15-10 for Tessera, k=2-49 for CellCharter, res=0.01-7.5 for STAGATE). For each method, we chose the resolution with the best average F1 score across all the benchmarked regions (**Fig S3b, S4a**).

## 7 SEA-AD MERFISH analysis

### 7.1 Data preprocessing

We downloaded raw MERFISH transcript files from the Seattle Alzheimer’s Disease Brain Cell Atlas (SEA-AD) data portal on AWS (https://registry.opendata.aws/allen-sea-ad-atlas/, accessed 3 May 2024). In total, the dataset included 75 middle temporal gyrus (MTG) samples from 27 donors. Transcripts were assigned to cells using the cell segmentation results from the original study^16^, which applied cellpose to the cytosolic and nuclear stains^40^. We annotated cell types with the Allen Institute’s MapMyCells tool (https://portal.brain-map.org/atlases-and-data/bkp/mapmycells), using the matching “10x Human MTG SEA-AD (CCN20230505)” reference taxonomy and the “SEA-AD Correlation Mapping” algorithm. As a result, each cell was labeled as belonging to one of 24 cell types identified in the corresponding SEA-AD snRNA-seq data. Cells were then filtered to keep only high quality cells with more than 30 transcripts and 15 genes per cell. The resulting spatial dataset had 1.7M cells, probed with a 140 gene panel.

We performed standard preprocessing of the single cell data using Seurat v5^41^. Transcript counts were log-normalized using the median number of transcripts per cell and Z-scored. Then the top 50 principal components (PCs) were computed using all features. We integrated cells across all 75 samples using Harmony v1.2, using the sample ID as the batch variable for correction^35^.

### 7.2 Tessera

We then constructed Tessera tiles using 50 PCs and minimum/maximum tile sizes of 5-10 cells (median 10 cells, IQR 8-14 cells). During Tessera, one round of gradient smoothing was performed using the “projected” distance and similarity metrics. We clustered Tessera tiles using Seurat’s FindNeighbors and FindClusters functions (with 50 PCs, 20 neighbors, and Louvain clustering with resolution 1.0), yielding 19 clusters. Tile clusters were annotated based on the cell types that were enriched in each.

### 7.3 Case-control analysis

We tested for associations between the abundance of each tile cluster and the continuous pseudo-progession score for each donor (CPS), a value between 0 and 1 previously calculated based on quantitative measurements of a number of pathological markers across the course of disease progression^16^. For each tile cluster, we fit a generalized linear model (GLM) using the `glm` function in R:

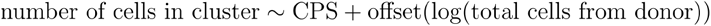

We modeled the count distribution using the quasipoisson distribution family to allow for overdispersion. When fitting the model, we aggregated multiple samples from the same donor, so each donor counted as a single independent observation. To avoid sampling bias in which different samples captured different amounts of cortex and white matter, we tested cells belonging to the cortex and white matter separately. For instance, we only considered cells belonging to one of the six white-matter oligodendrocyte-enriched clusters, excluding clusters that contain cortical layer neurons, or vice versa. Additionally, we tested for associations separately for early donors (12 individuals with CPS ≤ 0.4) and for late donors (15 individuals with CPS > 0.4), since prior work indicated distinct trajectories of disease progression in these groups^16^. To test for significant associations between each tile cluster and CPS, we performed a two-tailed T-test to test whether the coefficient of the CPS term was significantly nonzero. Finally, we performed multi-hypothesis test correction using the Benjamini-Hochberg method to control the false discovery rate^42^.

## 8. Reactive human lymph node Xenium 5K dataset

### 8.1 Data preprocessing

The complete FFPE Human Lymph Node with Xenium 5K Pan Tissue and Pathways Panel preview dataset was downloaded from the 10x Genomics website (https://www.10xgenomics.com/datasets/preview-data-xenium-prime-gene-expression). The sample consists of a single section of FFPE-preserved reactive left axillary lymph node. The gene expression panel targets 4,624 genes, with a total of 202,153,544 high-quality transcripts detected over an area of 48 mm^2^. We used the 708,983 cells that had been segmented using the 10x pipeline (based on nuclear, interior, and boundary stains), with a median of 255 transcripts assigned to each cell.

### 8.2 Cell type annotation

We annotated the cell types of individual cells in the Xenium 5K dataset using an external single-cell reference, which a previous study^17^ had compiled using scRNA-seq datasets from lymph nodes, spleen, and tonsils to capture a variety of immune cell states^18–20^. The scRNA-seq reference had 73,260 cells, with labels for 38 different cell types, including six germinal center (GC) B cell states, four non-germinal center (non-GC) B cell states, and fourteen T/NK/ILC cell states.

We performed reference label transfer using Harmony, annotating the Xenium 5K cells in two stages: first, we transferred coarse cell type labels, where the B and T/NK/ILC cell states were collapsed to single labels; then we transferred labels for the finer B and T/NK/ILC cell states separately for each of those subsets of cells.

For each stage of label transfer, only the genes that were shared between both Xenium 5K and scRNA-seq were used. Cell counts from both technologies were log-normalized to the same number of transcripts per cell (mean of median number of transcripts per cell in each technology). Features were then Z-scored, and weighted PCA used to compute the top 30 principal components. When computing weighted PCs, cells were inversely weighted by their representation across all cell types and studies, so that less frequent cell types still contributed to the PCs. The PCs were harmonized across technology, study, and sample ID variables using Harmony v1.2^35^. Single cells from the scRNA-seq reference and Xenium 5K were then jointly embedded by constructing a fuzzy 30-nearest neighbor connectivity graph using the UMAP algorithm^26,43^. Finally, reference labels were propagated along the connectivity graph until almost all the Xenium 5K cells were labeled. Note that some Xenium cells were too far away from or not in the same connected component as scRNA-seq cells. We left these cells unlabeled.

After coarse cell type annotation, finer cell type label transfer was done separately on these subsets of cells: GC B cells, non-GC B cells, and T/NK/ILC cells. The same Harmony-based label transfer procedure was used, except fewer genes and fewer weighted PCs were used (20 for GC B cells and T/NK/ILC cells, and 10 for non-GC B cells). The genes used for fine label transfer were selected by performing differential expression of the fine cell states on each subset of cells in the reference scRNA-seq dataset (yielding 740 significant genes for GC B cells, 386 for non-GC B cells, and 1,602 for T/NK/ILC cells).

### 8.3 Tessera

For the Xenium 5K single cells, we used the basic Seurat (v5) single-cell workflow to log-normalize the transcript counts, select the top 2000 highly variable genes, and Z-score the features^41^. The top 50 principal components (PCs) were computed using the highly variable genes. We then constructed Tessera tiles using 50 PCs and minimum/maximum tile sizes of 5-50 cells. During Tessera, one round of gradient smoothing was performed using the “projected” distance and similarity metrics.

We clustered Tessera tiles in two stages. First, we did coarse clustering of the aggregated tile embeddings using Seurat’s FindNeighbors and FindClusters functions (with 50 PCs, 20 neighbors, and Louvain clustering with resolution 1.0). We then performed subclustering of subsets of these original clusters to refine the labels for tiles enriched in GC B cells, non-GC B cells, T cells, and endothelium. When subclustering each subset of tiles, we recomputed principal components using the aggregated transcript counts for each tile, constructed a fuzzy nearest neighbor graph using the UMAP algorithm^26,43^, and performed Louvain clustering.

### 8.4 Downstream analysis

Tile clusters were annotated by calculating both the enrichment of specific cell types within each tile cluster and the colocalization of different tile clusters. Differential gene expression and differential cell type abundance were determined using a Wilcoxon rank sum test with presto^44^.

Individual lymphoid follicles were identified by performing Leiden clustering on the spatial tile adjacency graph, considering only tiles belonging to GC LZ/DZ, Mantle zone, cDC1/2, Macrophage/monocyte (2), or Non-GC cycling clusters. For each germinal center, the tiles belonging to the “Mantle zone” cluster were defined to form the mantle zone region, and the other tiles formed the germinal center (dark zone / light zone) region.

Germinal centers were then organized along a trajectory based on the cell type composition of the core dark/light zone region. For each germinal center, the fraction of cells belonging to each of 38 different annotated cell types was found. These features were mean-centered (but not scaled) and used to construct 10 PCs capturing major variation in cell type composition across the 41 different core germinal centers. A single germinal center trajectory was then computed using a principal curve (`princurve` package in `R`), with the 10 PCs as input features. Three of the smallest germinal centers (<10 cells) were excluded when constructing the trajectory to improve robustness. These germinal centers were then projected back onto the final trajectory after the trajectory was fit.

## 9. Lung IMC analysis

### 9.1 Data preprocessing

The complete lung cancer IMC dataset from Sorin et al^21^ was obtained upon request from the corresponding author. In total, the dataset included 1,633,610 cells from 416 patients with primary treatment-naive lung adenocarcinoma. Tissue microarrays were prepared from 1-mm^2^ cores from surgical tissue specimens.

Formalin-fixed paraffin-embedded tissue microarrays were profiled with 35-plex imaging mass cytometry (IMC). Per cell intensity was calculated across IMC channels using the cell segmentation results from the original study. Channel intensity was normalized by dividing by cell area.

### 9.2 Cell typing and QC

Low quality cells were filtered based on implausible (e.g. too large) morphology. Following extensive exploratory data analysis, we defined and removed morphological outliers with the following property: log(area) > 1.7 * log(area/perimeter) + 5. These preliminary “cells” all looked like long (200-1000um) and thin (<5um) rectangles along the edges of FOVs, likely arising from technical artifacts in segmentation. We also removed cells identified outside of tissue. We next robustly typed cells with our integrative annotation procedure, as follows: we normalized total expression of each channel by cell area, computed the top 16 PCs of the normalized feature-by-cell intensity matrix, and integrated over sample-specific effects in the PCs using Harmony^35^ v1.2. We then estimated a weighted nearest-neighbor graph of the cells in harmonized PC space and used the graph to estimate a two-dimensional UMAP embedding. We used the same graph to partition the cells by maximum modularity with the successive iterations of the Leiden algorithm, first with resolution parameter=1.2 and then resolution between 0.1 and 0.18 for successive subclustering. We next identified cluster markers with differential expression analysis. To do this, we used the Wilcoxon rank sum test as implemented in the presto package^44^. Cell-type labels were assigned to each cluster based on known functional state and lineage markers. Four clusters with insufficiently specific marker profiles were labeled “NA”. Finally, we performed sample level QC by filtering out samples that had more 20% NA cells. The resulting spatial data set had 1,493,455 cells across 385 samples profiled with a 35 protein panel.

### 9.3 Tessera and region identification

We constructed Tessera tiles using 10 harmonized PCs, minimum/maximum tile sizes of 5-20 cells, and otherwise default parameters. During Tessera, one round of gradient smoothing was performed using the “projected” distance and similarity metrics. We used Seurat’s FindNeighbors function to estimate a weighted nearest neighbor graph of the tiles in harmonized PC space and used the graph to estimate a two-dimensional UMAP embedding. We next used Seurat’s FindClusters function on the same graph to partition the cells by maximum modularity with the Louvain algorithm, yielding 17 clusters. Tile clusters were annotated as one of 10 different region types based on the cell types that were enriched in each.

### 9.4 Survival analyses

We performed Cox proportional hazards regression analysis to quantify the association between niche abundance and overall survival for each Tessera tile cluster (aka niche). Niche abundance was first normalized by calculating the per-sample proportion of area covered by each niche. We then used the coxph function from the survival package in R to construct a separate Cox proportional hazards model for each region type of the following form:

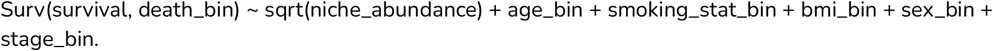

This formula models the hazard of death as a function of normalized niche abundance while adjusting for binarized age (age >=75), smoking status, binarized body mass index (BMI >=30), sex (M/F), and binarized disease stage status (stage I-II, stage III-IV) reported by the original authors. The coefficients for each niche feature, along with their corresponding confidence intervals and p values were extracted from the fitted models. To account for multiple testing, we adjusted the p-values using the Benjaminini-Hochberg method and reported survival associated clusters with a false discovery rate less than 20%.

We assessed differences in survival outcomes for stratification using multivariate Cox regression and LRT, similar to our strategy for testing survival association with Tessera region abundances. We used LRT (anova function in R stats package) to test the difference between the following null and alternative models, fit with the coxph function from the R survival package.

<u>**H0:**</u> Surv(survival, death_bin) ∼ age_bin + smoking_stat_bin + bmi_bin + sex_bin + stage_bin.

<u>**H1:**</u> Surv(survival, death_bin) ∼ stratification_group + age_bin + smoking_stat_bin + bmi_bin + sex_bin + stage_bin.

Above, stratification_group is a categorical, one-hot encoded variable denoting stratification group membership.

### 9.5 Co-abundance and stratification

We utilized the corr.test function from the psych package in R to calculate Spearman correlation coefficients and corresponding p-values between all pairs of normalized niche abundances. To account for multiple comparisons, we adjusted the p-values using the Holm–Bonferroni method to control the false discovery rate. We calculated the abundance of antitumor hubs by summing the per-sample proportion of area covered by the constituent regions. Patient stratification was performed by manual gating of the hub abundances per patient. Patients with Myeloid_Hub and Lymphoid_Hub abundances greater than or equal to 0.05 were reported as “Double-Positive.” Patients with Myeloid_Hub abundance greater than or equal to 0.05 and Lymphoid_Hub abundance less than 0.05 were classified as “Myeloid Hub Only.” Patients with Myeloid_Hub abundance less than 0.05 and Lymphoid_Hub abundance greater than or equal to 0.05 were classified as “Lymphoid Hub Only.” Patients with both Myeloid_Hub and Lymphoid_Hub abundances less than 0.05 were classified as “Pauci-Immune.”

## Data and Code Availability

All data used in analyses for this paper will be deposited into Zenodo prior to publication. All code to produce results and figures will be deposited into the github repository on korsunskylab/tesseramanuscript. The Tessera software is available as an R package on github.com/korsunskylab/tessera.

## Acknowledgements

This work was supported by the National Institutes of Health (5K01AR078355; NHGRI T32 HG002295) and the Chan Zuckerberg Foundation (Data Insights Grant). We thank Dr. Logan Walsh at McGill University for helping to access and process the lung cancer IMC data. We thank members of the Korsunsky, Raychaudhuri, and Hemberg labs at BWH for insightful discussions.

## Author contributions

DS and IK conceived the research. DS led computational work under the guidance of IK. MT led analysis of lung IMC data. All authors participated in interpretation and writing the manuscript.

## Figures

**Figure S1.**
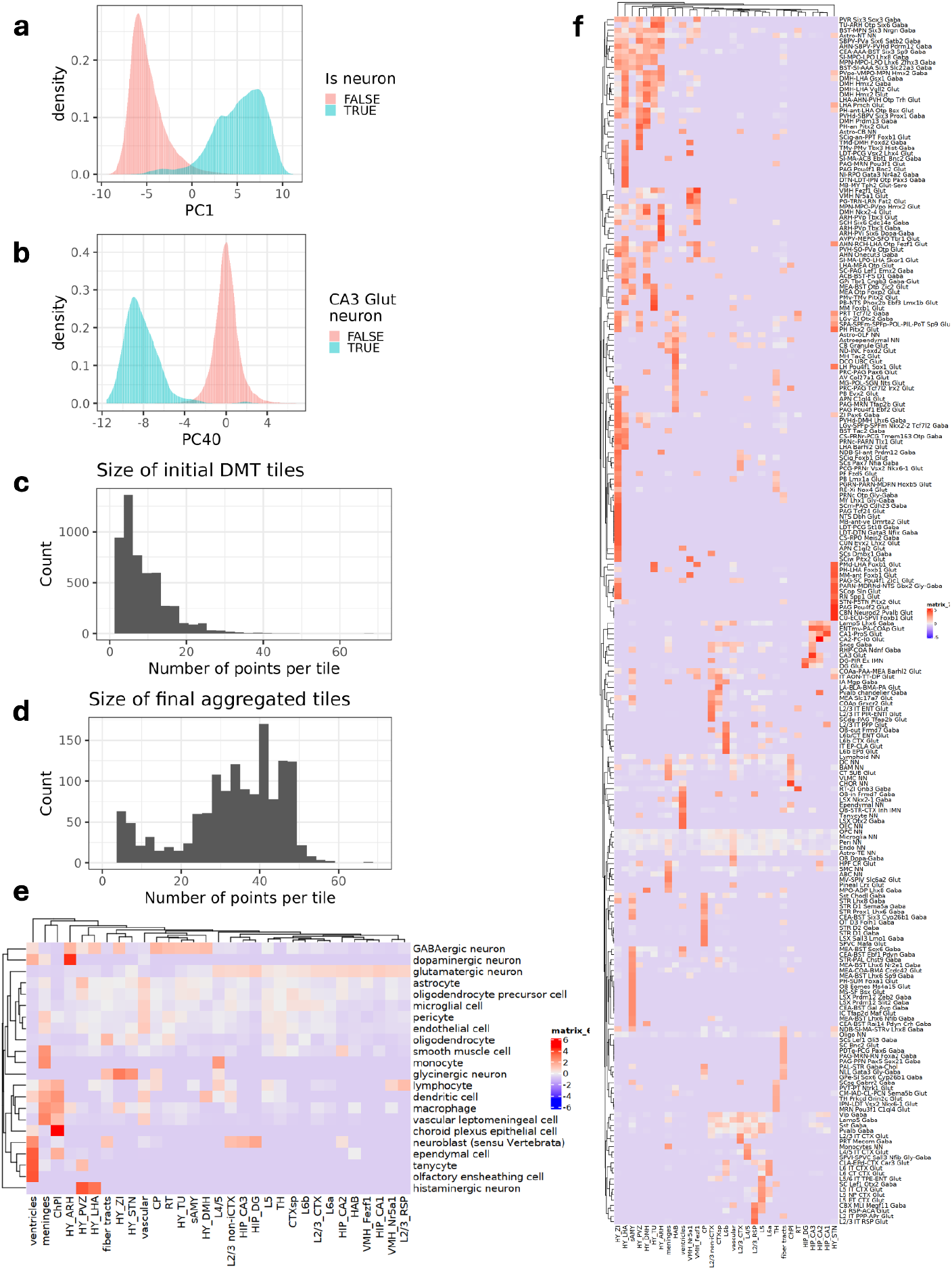
Tessera tile composition for a single section of the mouse brain. **a**, Distribution of PC1 embedding scores for neuronal and non-neuronal cells in the MERFISH data. **b**, Distribution of PC40 embedding scores for CA3 glutamatergic neurons and all other cells in the MERFISH data. **c**,**d**, Histogram of the number of cells per tile for initial tiles after topological segmentation (**c**) and final tiles after aggregation (**d**). **e**,**f**, Enrichment of cell types (**e**) and cell subclasses (**f**) in each Tessera tile cluster and sub-cluster.

**Figure S2.**
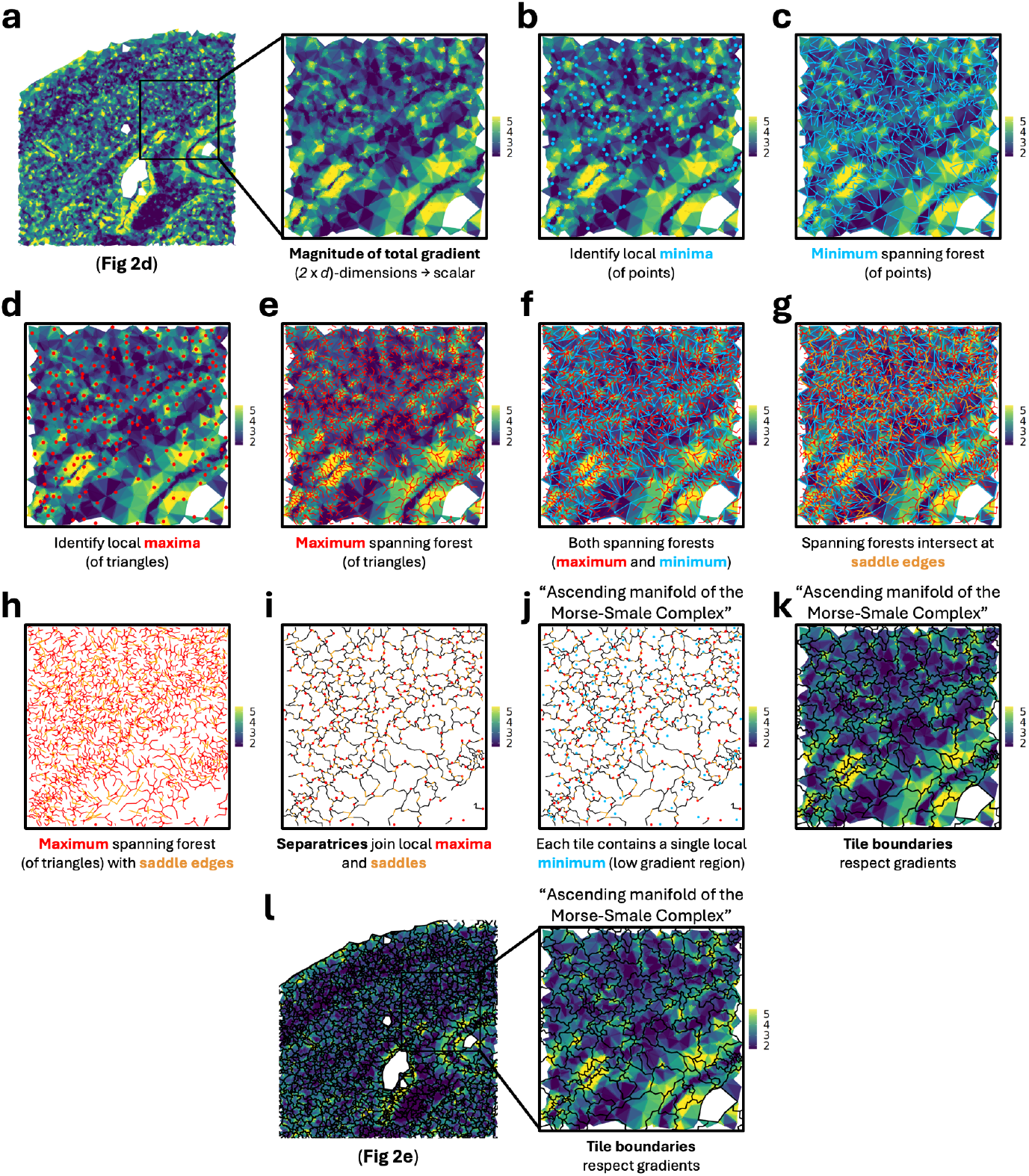
Topological segmentation using discrete Morse theory. Step-by-step example of a 900×900µm inset of the mouse brain section from (**Fig 2a-f**), demonstrating the initial construction of tile boundaries from the gradient field in (**Fig 2d-e**). **a**, Magnitude of the total gradient, with an inset of the region from (**Fig 2d**). **b**, Points (cells) that are local minima of the gradient field are shown in blue. **c**, Minimum spanning tree along the primal graph of points and edges (blue). **d**, Triangles that are local minima are identified in red. **e**, Maximum spanning tree along the dual graph of triangles and edges (red). **f**, Overlap of maximum and minimum spanning forests from (**c**) and (**e**). **g**, Shared saddle edges connecting trees in the minimum and maximum spanning forests are shown in orange. **h**, Maximum spanning forest (red) and saddle edges (orange) isolated from the overlay in (**g**). **i**, Separatrices (black) are defined as paths along the spanning forest connecting local maxima (red) with saddle edges (orange). **j**, Separatrices from (**i**) are plotted alongside the local minima from (**b**). **k**, Separatrices from (**i**) are overlaid on the gradient field from (**a**). The separatrices form the tile boundaries. **l**, Tile boundaries from topological segmentation, along with the underlying gradient magnitude, for the region in (**Fig 2e**). The inset matches (**k**).

**Figure S3.**
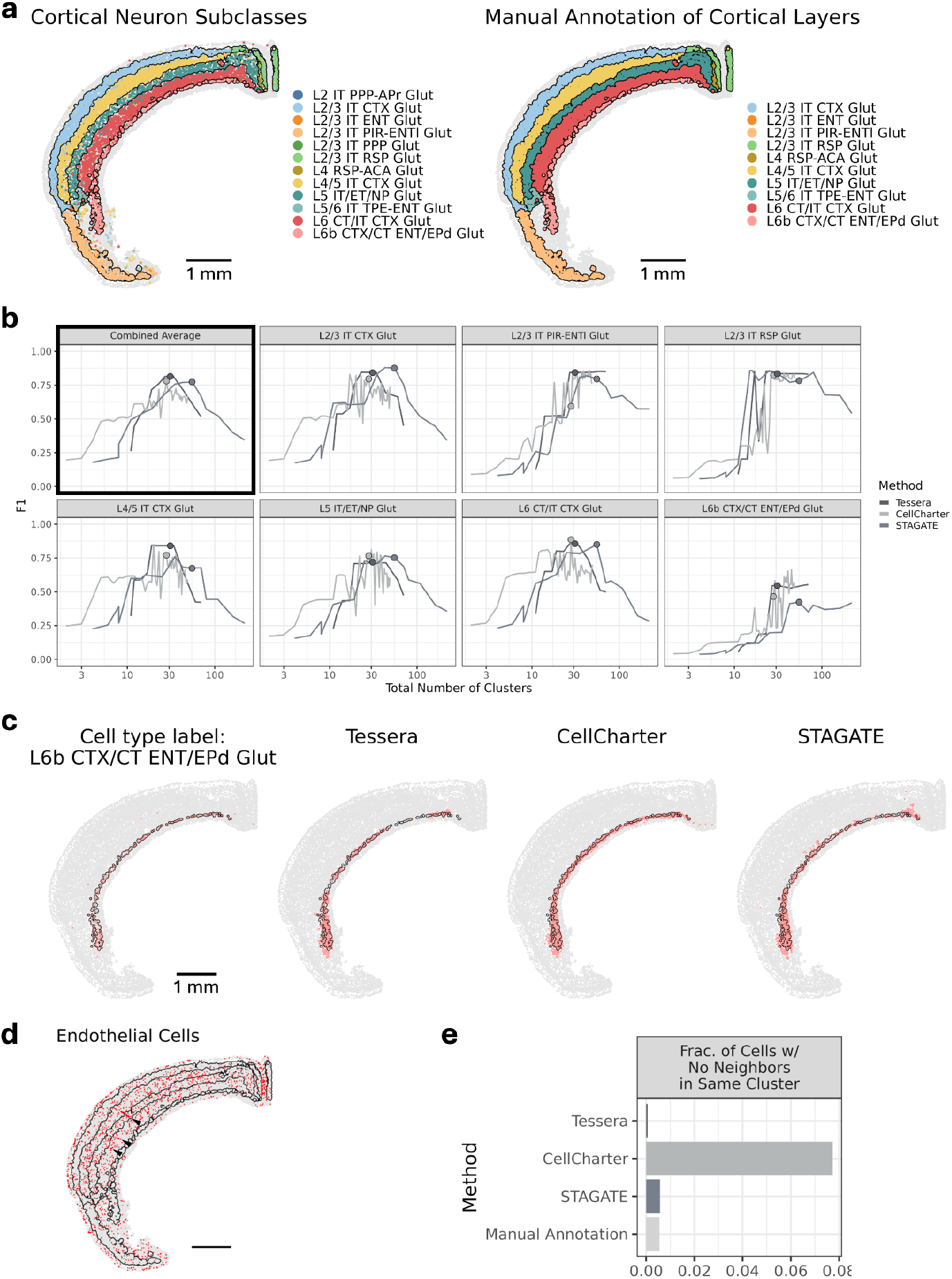
Benchmarking of Tessera, CellCharter, and STAGATE on a cortical layer segmentation task. **a**, Localization of labeled cortical layer-specific neurons (left) and manual region annotations (right). The same boundaries for the manually annotated regions are shown in both plots. **b**, F1-score for ventricle detection across a range of clustering resolutions. For each clustering result and region, the best matching cluster was evaluated. The combined average F1-score is a weighted average of the F1-score for each region, weighted by the size of the region. For each method, the clustering resolution that maximized the combined average F1-score is highlighted (Tessera, Leiden resolution = 2, with 31 clusters; CellCharter, GMM k = 28, with 28 clusters; STAGATE, Leiden resolution = 1, with 55 clusters). These are the resolutions shown in (**Fig 3b**). **c**, Cell type labels and spatial domains matching cortical layer L6b, using the three methods Tessera, CellCharter, and STAGATE, with the same clustering resolutions as in (**Fig 3b**) (Leiden resolution = 2, GMM k = 28, and Leiden resolution = 1, respectively). **d**, Individual endothelial cells (red) within the cortical regions. Black boundaries match the manually annotated regions in (**a**). Arrowheads identify clusters of endothelial cells. **e**, Fraction of cells within the cortical region that are singletons, with no neighbors assigned to the same cluster.

**Figure S4.**
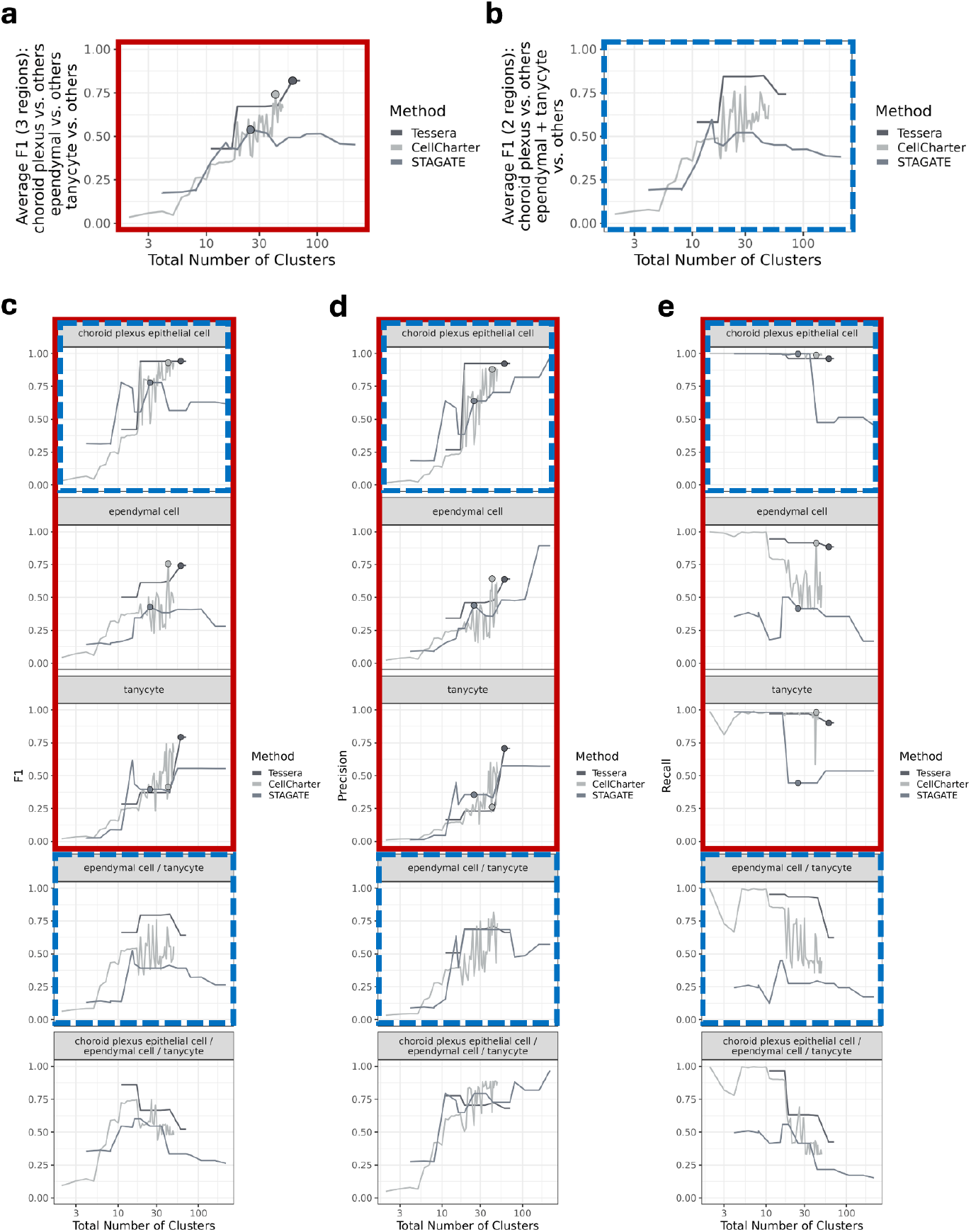
Benchmarking of Tessera, CellCharter, and STAGATE on a ventricle segmentation task. **a**, Average F1-score for detecting 3 distinct ventricle regions—choroid plexus, ependyma, and tanycyte (average of red regions in (**c**))—across a range of clustering resolutions. For each clustering result and region, the best matching cluster was evaluated. The combined average F1-score is a weighted average of the F1-score for each region, weighted by the size of the region. For each method, the clustering resolution that maximized the average F1-score is highlighted (Tessera, Leiden resolution = 7.5, with 60 clusters; CellCharter, GMM k = 42, with 42 clusters; and STAGATE, Leiden resolution = 0.3, with 25 clusters). These are the resolutions shown in (**Fig 3f**). **b**, Average F1-score for detecting 2 distinct ventricle regions—choroid plexus and combined ependyma/tanycyte (average of blue regions in (**c**))—across a range of clustering resolutions. **c-e**, F1-score, precision, and recall for detecting 5 distinct ventricle regions: 1) choroid plexus, 2) ependyma, 3) tanycyte, 4) combined ependyma/tanycyte, and 5) combined choroid plexus/ependyma/tanycyte (all ventricle regions). Results are shown across a range of clustering resolutions, with the best overall resolution from (**a**) highlighted (Tessera, Leiden resolution = 7.5, with 60 clusters; CellCharter, GMM k = 42, with 42 clusters; and STAGATE, Leiden resolution = 0.3, with 25 clusters).

**Figure S5.**
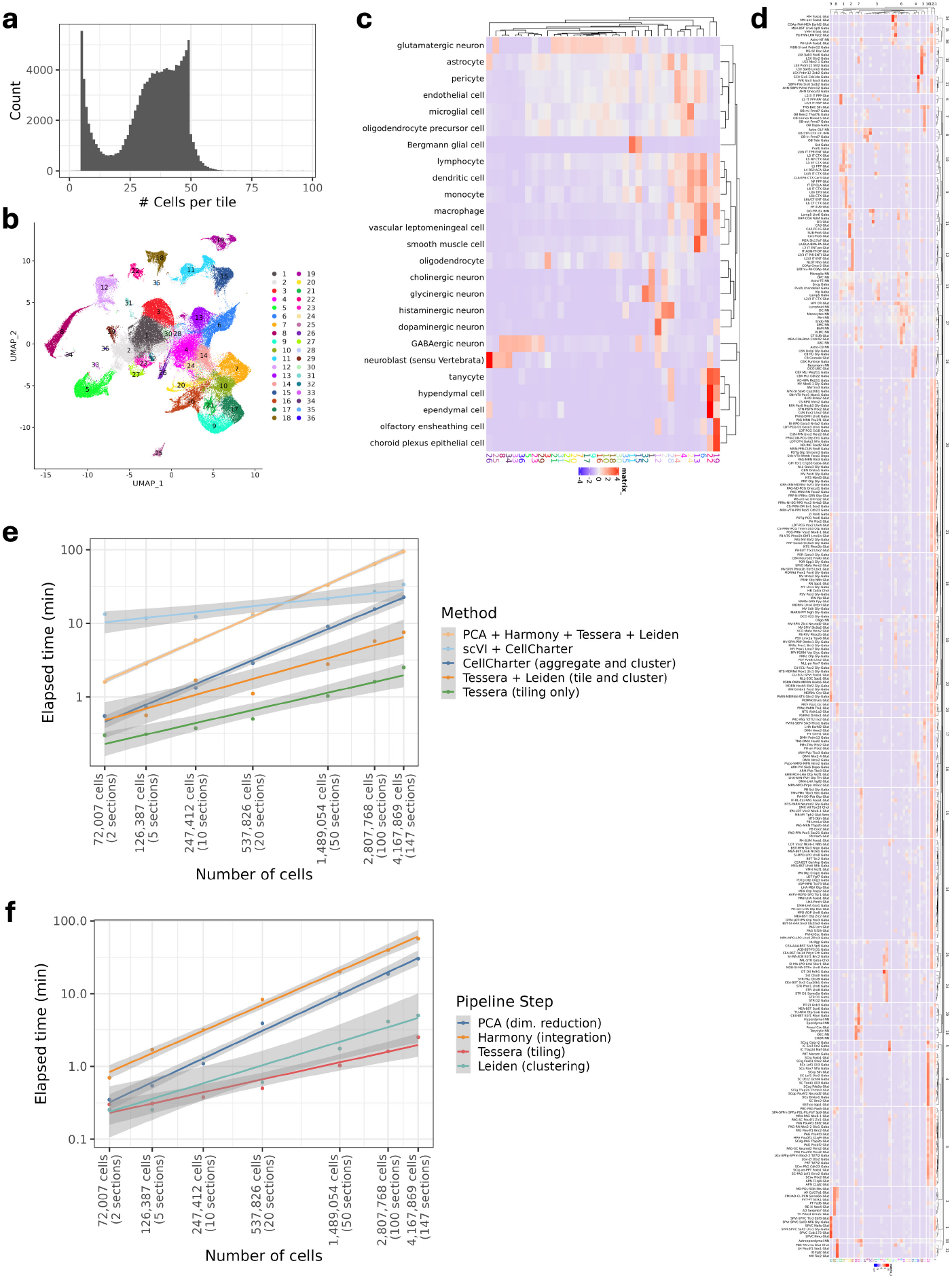
Scalability of Tessera across 147 sections and 4.2 million cells spanning the whole mouse brain. **a-d**, Additional Tessera results from the analysis in (**Fig 4**), where Tessera was applied to all 147 sections across the whole mouse brain. **a**, Histogram of the number of cells in each tile. **b**, UMAP visualization, where each point represents a tile, showing 36 clusters obtained using Leiden clustering. The colors match the tile clusters in (**Fig 4**). **c**,**d**, Enrichment of broad cell types (**c**) and cell subclasses (**d**) within each tile cluster. Red (blue) indicates a cell type is found within a tile cluster more (respectively, less) often than expected by chance if cells were uniformly distributed. **e**, Time required for different stages of the Tessera pipeline (with 20-core CPU) and CellCharter pipeline (with GPU), applied to subsets of sections ranging from 2 sections (72K cells) to 147 sections (4.2M cells). Dimensionality reduction and integration are typically pre-computed, so performance is reported both with and without these steps. **f**, Time required for each step of the Tessera analysis pipeline, run with 20 CPUs.

**Figure S6.**
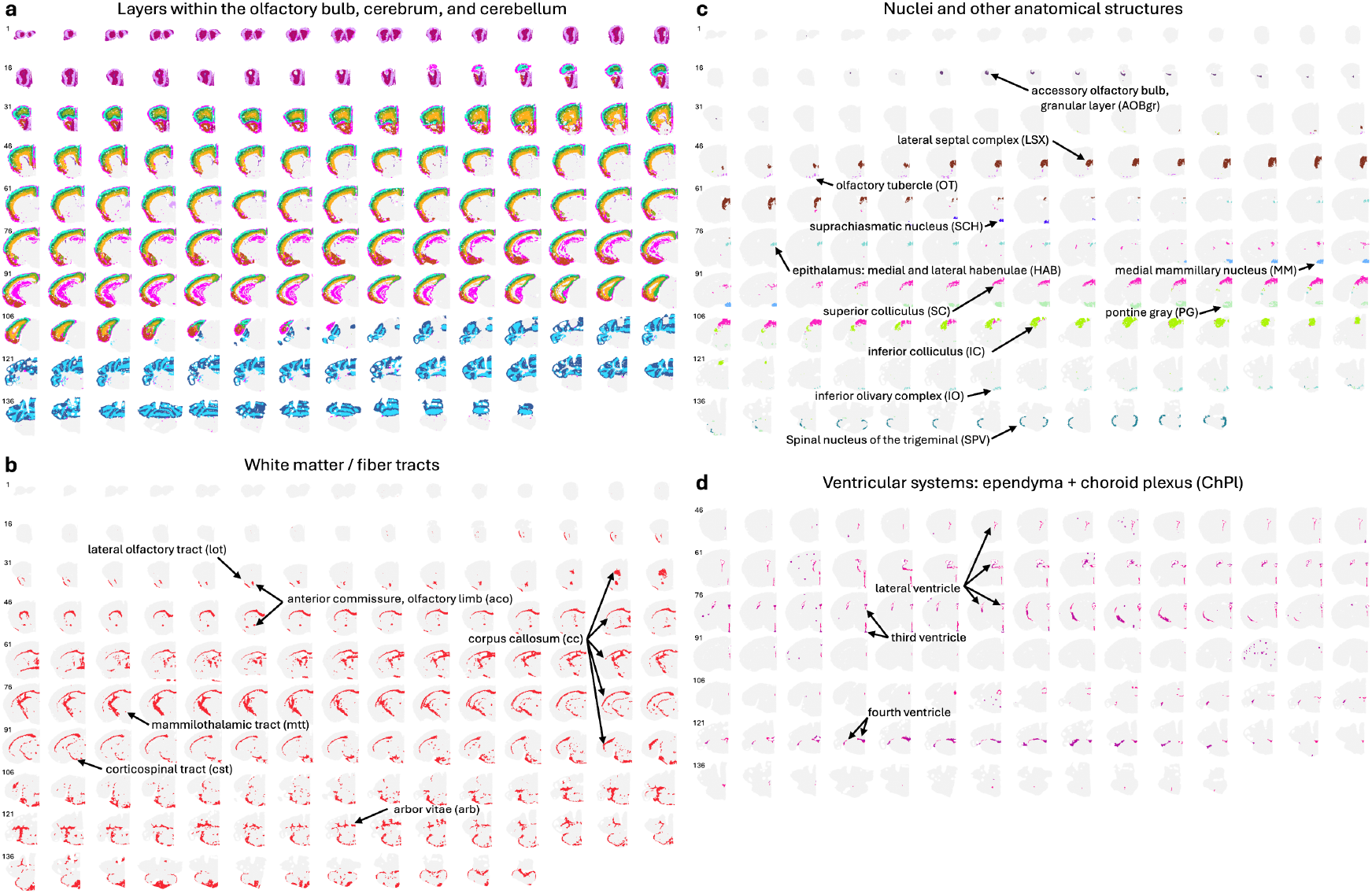
Tessera tiles correspond to true anatomical structures spanning the whole mouse brain. **a-d**, Spatial localization of subsets of Tessera tile clusters from the analysis in (**Fig 4**), where Tessera was applied to all 147 sections across the whole mouse brain. Sections are ordered from anterior to posterior, and the colors match the tile clusters in (**Fig 4**).

**Figure S7.**
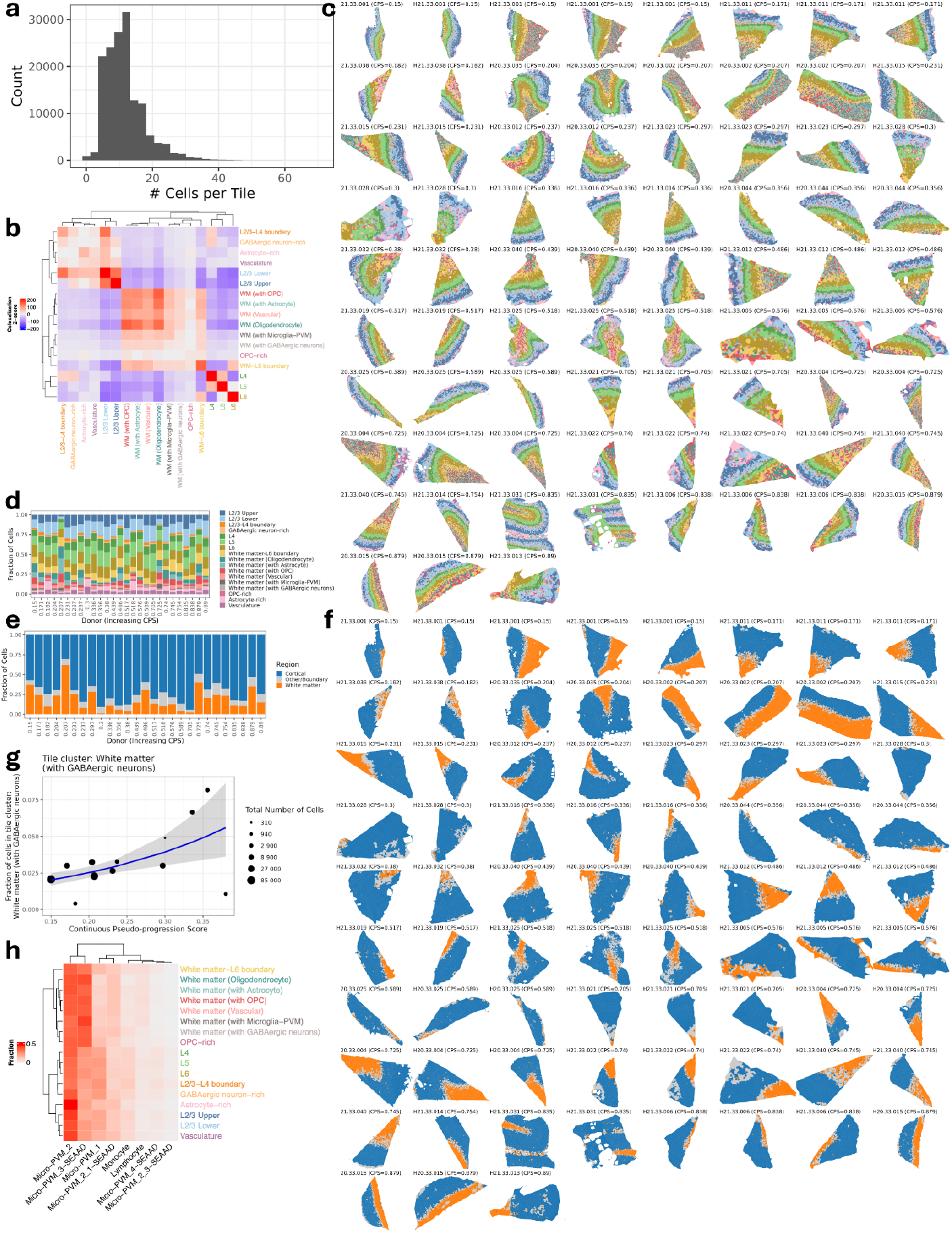
Characterizing Tessera tile clusters from the Seattle Alzheimer’s Disease Brain Cell Atlas (SEA-AD) dataset. Tessera tiles correspond to the analysis from (**Fig 5**). **a**, Histogram of the number of cells in each tile. **b**, Colocalization of tile clusters across all samples. Red (blue) indicates a tile cluster is adjacent to another tile cluster more (respectively, less) often than expected by chance if tile cluster labels were randomly permuted. **c**, Spatial localization of tile clusters across all samples (75 samples from 27 donors). The colors match the tile clusters in (**Fig 5a**). **d**, Fraction of cells belonging to each tile cluster for each donor, arranged by increasing CPS. **e**, Fraction of cells belonging to cortical and white matter regions for each donor, arranged by increasing CPS. **f**, Spatial localization of cortical and white matter regions across all samples. These regions are defined by combining the corresponding tile clusters and filling any holes. **g**, Association between the fraction of white matter cells in the GABAergic neuron-enriched white matter tiles and CPS for early donors (CPS ≤ 0.4). Each point represents samples from a single donor, where the point size corresponds to the total number of white matter cells from that donor. **h**, Fraction of microglia-PVM cells (between 0 and 1) belonging to each supertype within each tile cluster. Rows are normalized to sum to one.

**Figure S8.**
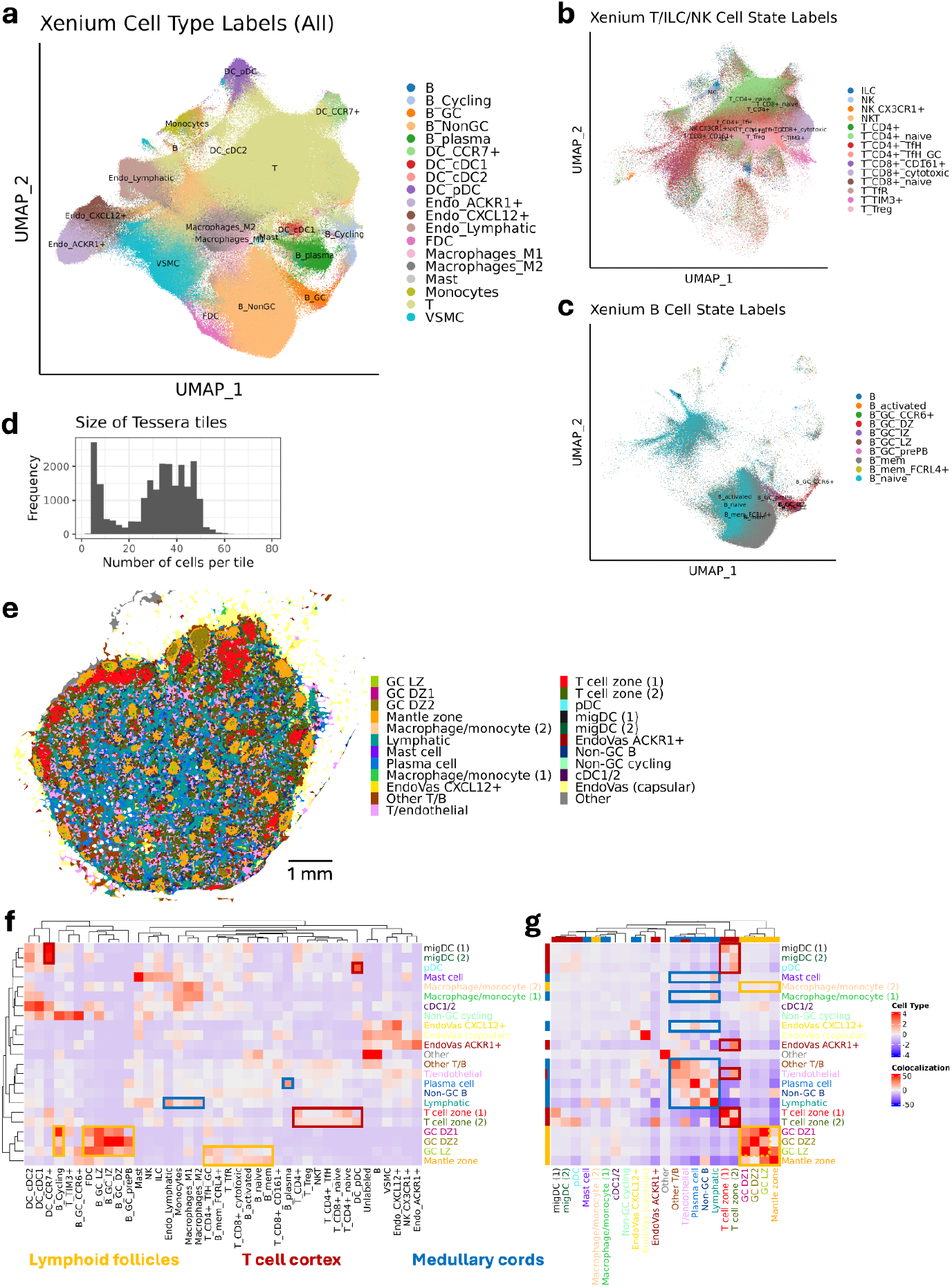
Annotation of single cells and Tessera tiles from Xenium 5K profiling of a lymph node. **a-c**, UMAP visualization of single cells from Xenium 5K profiling, colored by cell type labels after annotation using an external scRNA-seq reference (**Methods**). Finer cell state labels are shown for the T and B subsets of the lymphocyte populations (**b**,**c**). **d**, Distribution for the number of transcripts per single cell from Xenium. **e**, Spatial localization of all tile clusters in (**Fig 6a-e**). **f**, Cell type enrichment within each Tessera tile cluster. Positive values indicate a cell type is observed in a given tile cluster more frequently than expected by chance if cells were randomly distributed among tiles. **g**, Colocalization of tile clusters. Positive values indicate tiles belonging to two given clusters are adjacent to each other more frequently than expected by chance if cluster labels were permuted.

**Figure S9.**
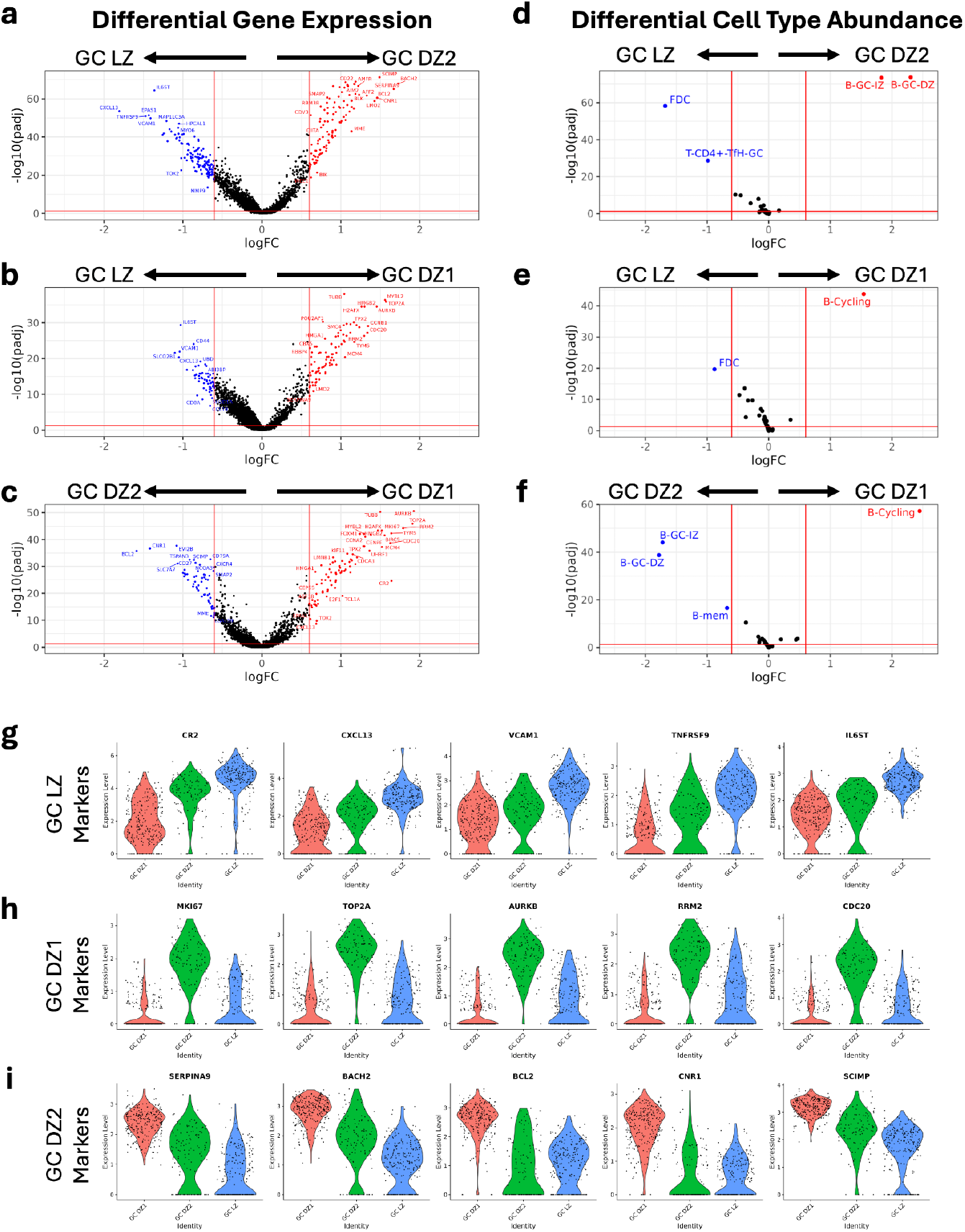
Differential expression between germinal center tile clusters. **a-c**, Pairwise differential gene expression between three germinal center (GC) tile clusters—GC LZ, GC DZ1, and GC DZ2 (**Fig 6a**). Differential expression analysis was performed using a Wilcoxon rank sum test. **d-f**, Pairwise differential cell type abundance between the three GC tile clusters. Differential abundance analysis was performed using a Wilcoxon rank sum test. **g-i**, Violin plots for expression of the top gene markers in the GC tile clusters.

**Figure S10.**
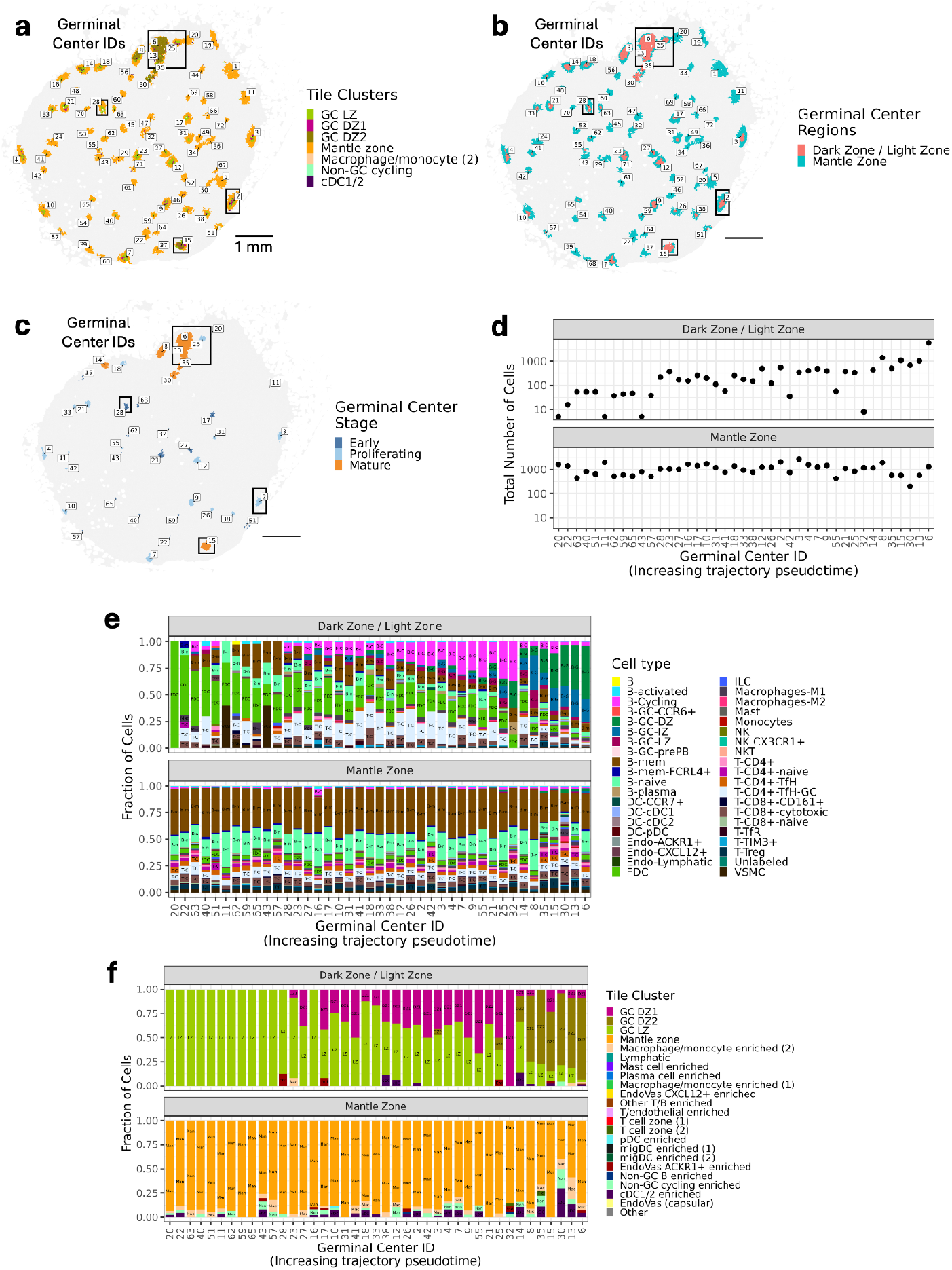
Definition and analysis of individual germinal centers. **a**, Identification of individual germinal centers based on colocalized tile clusters that compose the germinal centers. **b**, Definition of germinal center (GC LZ/DZ1/DZ2 tile clusters) and mantle zone (mantle zone tile clusters) regions of the germinal centers. **c**, Spatial localization of individual germinal centers belonging to different stages of the trajectory in (**Fig 6l**). **d**, Size of the dark/light zone core and mantle zone periphery regions of individual germinal centers. **e**,**f**, Cell type (**e**) and tile cluster composition (**f**) of the dark/light zone and mantle zone regions for all 41 germinal centers, ordered according to the trajectory in (**Fig 6l**).

**Figure S11.**
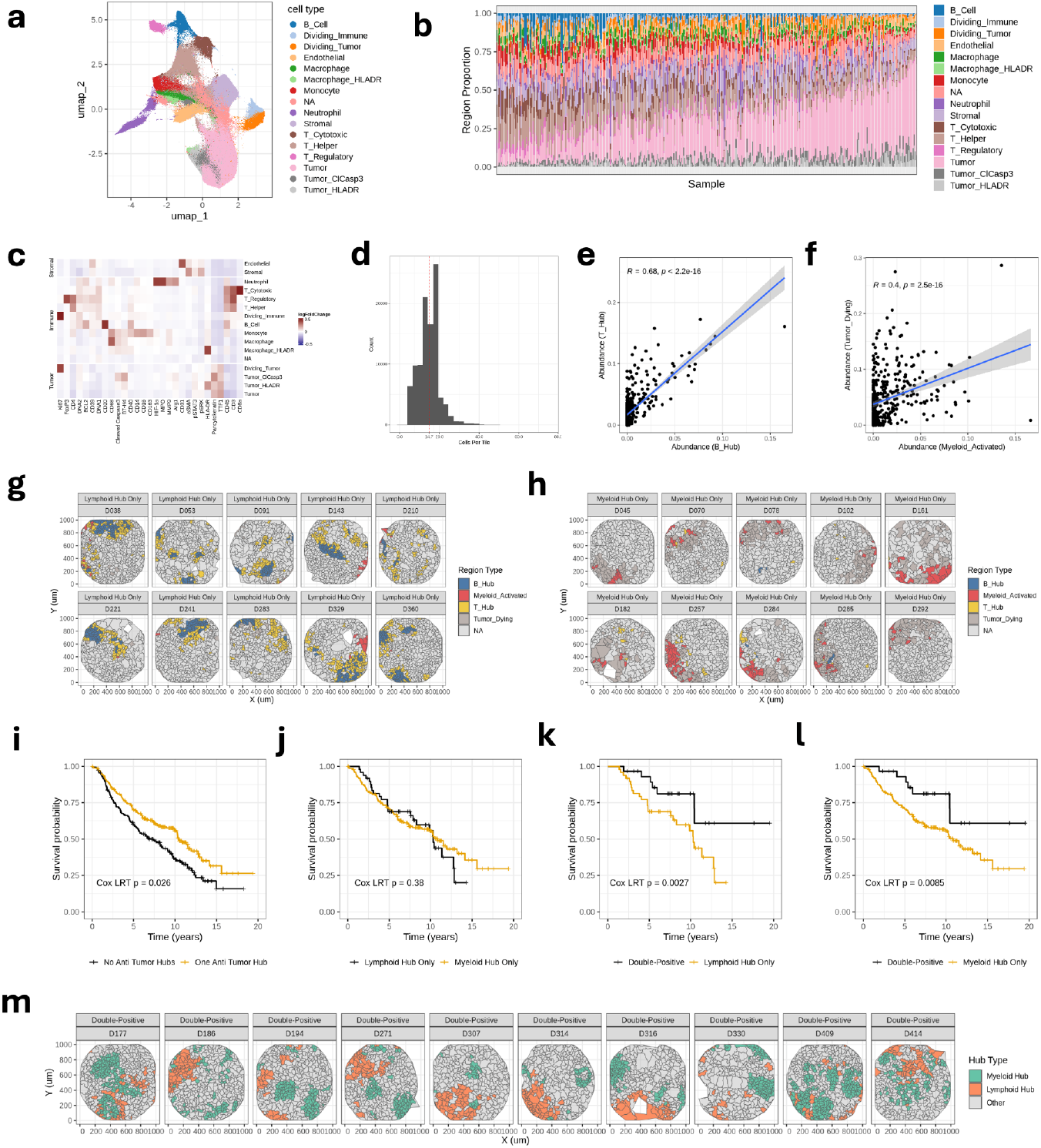
Annotation of single cells, characterization of Tessera tiles, and survival analyses from IMC profiling of 385 LUAD patients. **a**, UMAP projection of cells, labeled by integrated cell type annotations. **b**, Proportion of each cell type per sample. **c**, Differential expression statistics (log2 fold change) for protein markers in each cell type. **d**, Histogram of number of cells per Tessera tile. Red line denotes the median. **e**, Correlation of the proportion of B_hub region per sample (x-axis) with proportion of T_hub region per sample (y-axis). R and p-value from Spearman correlation test. **f**, Correlation of the proportion of Myeloid_Activated region per sample (x-axis) with proportion of Tumor_Dying region per sample (y-axis). R and p-value from Spearman correlation test. **g**,**h**, Representative examples of samples with lymphoid (**g**) and myeloid (**h**) anti-tumor hubs, highlighting the four region types that define these hubs. **i-l**, Two-group survival distribution comparisons, with multivariate Cox regression p-values and Kaplan Meier survival curves, comparing outcome of (**i**) pauci-immune group (black) to combined lymphoid-only and myeloid-only groups (orange), (**j**) lymphoid-only (black) and myeloid-only (orange) groups, (**k**) double-positive group (black) to lymphoid-only group (orange), and (**l**) double-positive group (black) to myeloid-only group (orange). **m**, Representative spatial plot of samples in the double-positive subgroup, highlighting the colocalization of lymphoid and myeloid anti-tumor hubs.

